# Linking Neural Manifolds to Circuit Structure in Recurrent Networks

**DOI:** 10.1101/2024.02.28.582565

**Authors:** Louis Pezon, Valentin Schmutz, Wulfram Gerstner

## Abstract

The classic view of cortical circuits composed of precisely tuned neurons hardly accounts for large-scale recordings indicating that neuronal populations are heterogeneous and exhibit activity patterns evolving on low-dimensional manifolds. Using a modelling approach, we connect these two contrasting views. Our recurrent spiking network models explicitly link the circuit structure with the low-dimensional dynamics of the population activity. Importantly, we show that different circuit models can lead to equivalent low-dimensional dynamics. Nevertheless, we design a method for retrieving the circuit structure from large-scale recordings and test it on simulated data. Our approach not only unifies cortical circuit models with established models of collective neuronal dynamics, but also paves the way for identifying elements of circuit structure from large-scale experimental recordings.

## Introduction

The progress of experimental methods continuously reshapes our understanding of the link between brain activity and behaviour. While, for a long time, neuroscientists could only record one or a few neurons at the same time, the recent availability of tools such as high-density probes and two-photon calcium imaging has enabled the simultaneous recording of thousands of neurons [1]. As a consequence, systems neuroscience currently offers two complementary perspectives on the structure of neuronal activity [2, 3].

The first perspective focuses on the representation and processing of information by single neurons characterised by individual functional properties. As a starting point, landmark studies in neuroscience have established that the activity of single neurons can be modulated by features of sensory stimuli [4–6], spatial location and orientation in the environment [7–11], or higher-level task-related variables [12–14]. The dependence of single-neuron responses to stimulus- or task-related features, usually termed *tuning*, can generally be summarised by a few properties, such as the centre of their receptive field. For example, in flies and mammals, the sense of spatial orientation arises from a population of head-direction cells, where each cell responds preferentially to a specific head direction (HD) [9, 10, 15]. Cells can thus be sorted according to their functional role in the head direction circuit by placing them on a ring (Fig. 1A), where neighbouring cells have similar preferred HD [16, 17]. Single-neuron properties might also describe other response features besides stimulus tuning: neuronal responses can be modulated by some task- or stimulus-independent variables that are not controlled by the experimenter, such as internal brain state [18–22]. In this view, each neuron can be assigned a “location”, i.e. be embedded, in an abstract space where neighbourhood reflects the functional similarity between neurons. We call it the *functional similarity space*, or more briefly *similarity space*. Importantly, the embedding of the neurons may not correspond to their physical location in the brain [22]. For instance in mammalian V1, preferred orientations are distributed across the cortical sheet, forming pinwheel-like patterns [23, 24].

**Fig. 1.**
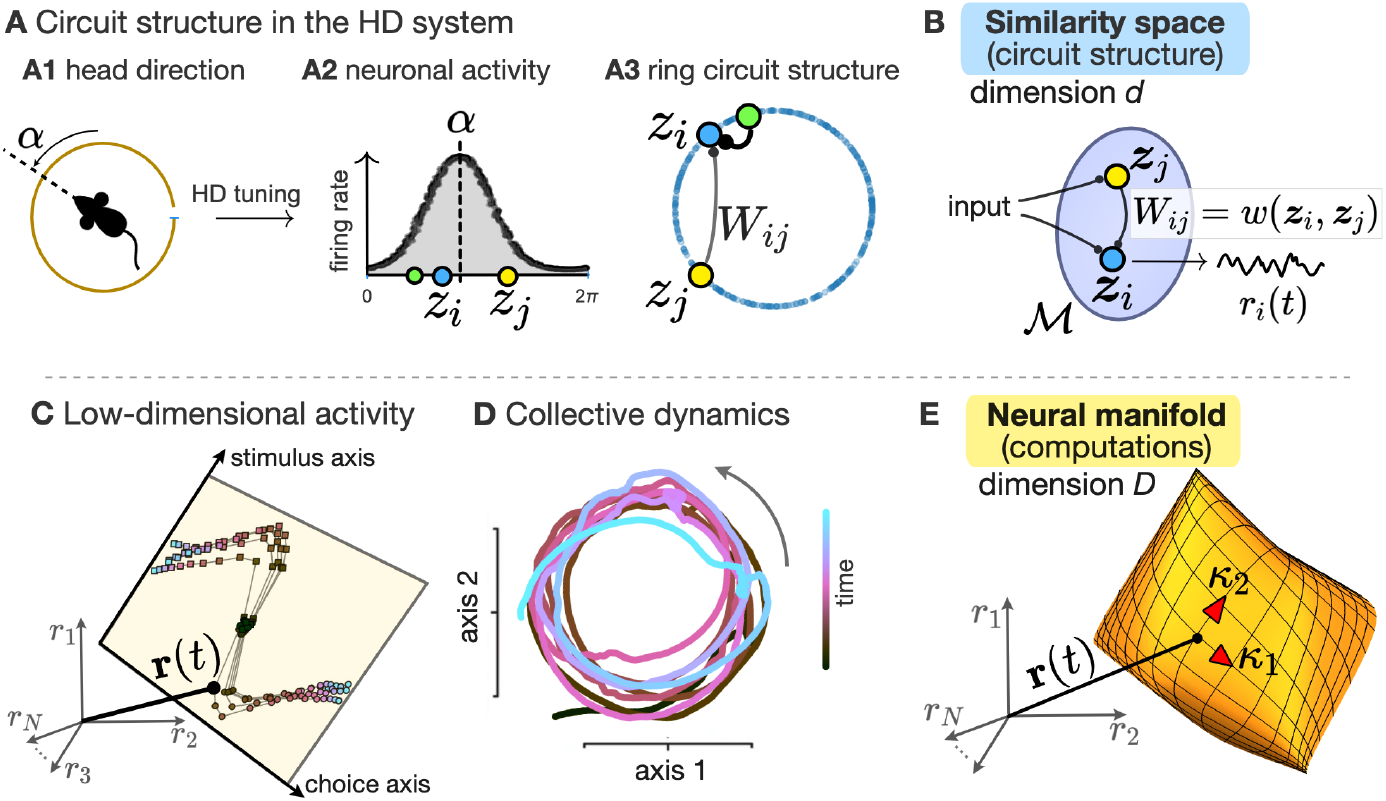
Two perspectives on neuronal activity. **A**. The “circuit” perspective illustrated by the ring model of the head direction (HD) system. **A1**. The animal’s HD is represented by an angle α. **A2**. Each neuron *i* in the population responds preferentially to a specific HD angle *z*_*i*_ (horizontal axis). Neuronal activity (vertical axis) depends on the difference between the current HD α and each neuron’s preferred HD angle *z*_*i*_. Coloured dots represent three examples neurons. **A3**. Each neuron’s preferred HD angle determines its “location” on a one-dimensional ring (blue dots). The persistent activity of HD cells in the absence of stimulation [42, 46] can be explained by a ring attractor model [16, 17], where recurrent synaptic connections *W*_*ij*_ between pairs of neurons *i* and *j* are stronger if neurons are close neighbours on the ring (e.g. the blue and green neuron) than if they are further apart (e.g. the blue and yellow neuron). **B**. In the “circuit” perspective, single-neuron responses *r*_*i*_(*t*) are characterised by the locations of the neurons in an abstract *similarity space* ℳ of dimension *d*, where neighbourhood represents functional similarity. Classical circuit models postulate that recurrent connections between pairs of neurons *i* and *j* (and forward connections from external inputs) are determined by their locations ***z***_*i*_, ***z***_*j*_ in this abstract space. **C**. The “neural manifold” perspective illustrated by large-scale recordings of the neuronal activity in the monkey prefrontal cortex [34]. During a decision-making task, the population activity **r**(*t*) of a population of *N* cells lives in a two-dimensional subspace, spanned by a ‘stimulus’ and a ‘choice’ axis, which encode information about the sensory stimulus and the decision made during a trial, respectively. Connected points correspond to the population activity during a single trial, with color representing time elapsed since trial onset. **D**. In mouse medial entorhinal cortex, the linear projection of the population activity onto relevant axes reveals slow oscillatory population-level dynamics, internally generated by recurrent mechanisms [22]. **E**. In the “manifold” perspective, the joint activity of *N* neurons, **r**(*t*) ∈ ℝ^*N*^, lies on a low-dimensional *neural manifold* with intrinsic dimension *D* ≪ *N* (shown here as a curved, (*D* = 2)-dimensional surface). The activity is parametrised by population-level latent variables *κ*_1_(*t*), …, *κ*_*D*_(*t*) whose dynamics reflect neural computations. Panel (C) is inspired from [20]. Panel (D) is inspired from [22].

The second perspective takes a descriptive, population-level approach, put forward by the advent of large-scale recordings [25]. In this perspective, the joint activity of *N* simultaneously recorded neurons, or *population activity*, is represented at any given time as a point in a *N*-dimensional state-space. Several studies have revealed that the activity of heterogeneous, large populations of neurons is often restricted to a *low-dimensional* space, often referred to as the “neural manifold”, embedded within the *high-dimensional* state-space of the population activity [26–31]. In the neural manifold, the momentary activity can be described by a small number *D* ≪ *N* of time-dependent variables that represent computationally relevant quantities; e.g., the animal’s head direction, or the perceived sensory stimulus (Fig. 1C). The change of these variables over time reveals the low-dimensional dynamics of the population activity on a time scale from tens of milliseconds [21, 32–34] to tens of seconds [22] (Fig. 1D). Hence, in this view, neural computation is described by a set of trajectories, or “flow”, in the low-dimensional neural manifold (Fig. 1E) that emerges from the collective dynamics of a large number of neurons [34–38].

To summarise, the first perspective focuses on how the functional properties of individual neurons shape neuronal activity, whereas the second perspective focuses on a population-level description of neural representations and computations [2]. The concept of tuning potentially provides a direct link between the two perspectives [39]. A tuning curve traditionally describes how a single neuron’s response depends on a particular stimulus variable. Thus, the set of tuning curves of all the neurons in the population directly maps each possible stimulus to the corresponding population activity. If the set of all possible stimuli is parameterised, say, by a one-dimensional circular variable (for example, the head direction in the case of HD cells), the set of all the possible population activities will in turn necessarily form a ring. More generally, low-dimensional tasks tend to yield low-dimensional neural trajectories [40, 41]. Yet, such a static transformation from stimulus to population activity is insufficient to explain some of the most striking features of neural manifolds. First, it has been observed that neural manifolds can persist during sleep [27, 28, 42], and be dissociated from sensory cues [10, 43]; which implies that the neural manifold is not a mere reflection of external stimuli. Second, the low-dimensional trajectory of the population activity observed in experiments often does not correspond to the evolution of present external stimulus variables, but is rather determined by the initial condition [21], context [34], or internally-generated factors [22, 30, 44] (Fig. 1D). Both findings point towards underlying recurrent circuit mechanisms [26, 45].

In the examples mentioned above, strong excitatory connections are found between neurons with similar preferred HD in the fly HD system [47], or between neurons with a similar orientation tuning in V1 [24]. Consistent with the traditional notion of cortical columns [48], these observations support the idea that functional similarities between neurons arise from network structure [12, 42, 49]. This idea has been the basis for many computational models in which forward and recurrent connections are determined by the neurons’ respective functional properties (Fig. 1B) [50–57]. However, the link between so-called *circuit models* and neural manifolds remains elusive [2, 3].

In this paper, our aim is to connect the two perspectives by addressing the following questions.

i. Can we establish a mathematical link between the similarity space, summarising the structure of the connectivity in circuit models, and the flow of the collective dynamics in the neural manifold?
ii. Is there a one-to-one mapping between the neural manifold and the similarity space? Although the topology of the neural manifold has been found to match that of the similarity space in special cases (for example, a ring in the fly HD system) [3, 47], there is no obvious reason why this should be true in general. (iii) Can we extract knowledge about the circuit structure from neuronal activity? Although many studies have focused on extracting the neural manifold from experimental data [27, 28, 32, 33], an established method for extracting the similarity space, without explicitly referring to tuning curves or selectivity, is missing.

The results are organised as follows. First and foremost, we propose a modelling framework that provides an explicit link between circuit structure and collective dynamics in Recurrent Neural Networks (RNNs). Relying on the theory of neural fields [52] and large low-rank networks [58], our framework connects functional properties of single neurons to emergent low-dimensional dynamics in an RNN model. Maybe surprisingly, the theory predicts that there is no one-to-one relationship between the functional similarity space and the neural manifold. In particular, their respective dimensionality can be different, and the same collective dynamics can result from different circuit structures. Finally, we propose a method for extracting the similarity space from the neuronal activity, by exploiting the similarities between the activity of single neurons. We apply this method to simulated data, and show that we can retrieve the dissimilar circuit structure of different RNNs performing the same simple task.

## Results

### A unifying framework based on the theory of large RNNs

In this section, we present a modelling framework that links the *similarity space* of single-neuron response properties with the *neural manifold* of population activity; the details of the derivations are presented in the Methods 1. Our framework extends recent theoretical results on large RNNs [59–61] to field models [52, 59]. Since the dynamics of large Spiking Neural Networks (SNNs) composed of Linear-Nonlinear-Poisson (LNP) neurons [62, 63] converges to the dynamics of equally large RNNs in the large *N* limit (with 1/*N* synaptic weight scaling) [61], we directly consider the case of LNP spiking neurons.

In a network composed of *N* neurons, let the membrane potential *h*_*i*_ of each neuron *i* = 1, …, *N* be a leaky integrator (with time constant *τ*) of recurrent and external inputs. Each spike of a presynaptic neuron *j*, connected to postsynaptic neuron *i*, causes an instantaneous change of the postsynaptic potential *h*_*i*_ with magnitude given by the synaptic weight 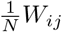 from neuron *j* to neuron *i*. The network also receives *K* external inputs *I*_1_(*t*), …, *I*_*K*_(*t*), with input weights to neuron *i* denoted by 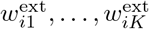, respectively. The dynamics of the membrane potential *h*_*i*_ thus reads:

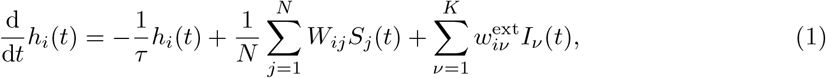

where 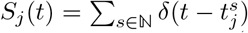 denotes the spike train of neuron *j*. The instantaneous firing rate of each neuron *i* is given by *r*_*i*_(*t*) = *ϕ*(*h*_*i*_(*t*)), where *ϕ* is a positive, *nonlinear* transfer function. The set of instantaneous firing rates, **r**(*t*) = (*r*_1_(*t*), …, *r*_*N*_ (*t*)), define the intensity of *N* (conditionally independent) inhomogenous *Poisson* processes from which the spike times 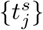 of each neuron *j* are drawn.

The theoretical framework relies on two assumptions. First, we assume that each neuron *i* can be functionally characterised by a small set of *d* ≪ *N* features, denoted ***z***_*i*_ = (*z*_*i*,1_, …, *z*_*i,d*_), which determine its abstract location in a *d*-dimensional *functional similarity space* ℳ, where neurons close to each other have similar response properties. This assumption is rooted in the circuit perspective described in the introduction and is the starting point of many models in computational neuroscience [16, 50–57]. This assumption means that both recurrent and external connections in Eq.(1) depend on the respective “locations” of the neurons, so that Eq.(1) can be rewritten as:

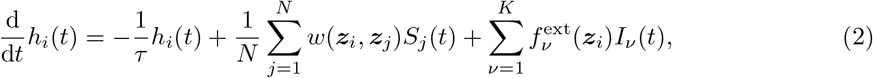

where 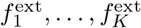 are continuous functions that characterise the tuning of the neurons to the external inputs *I*_1_, …, *I*_*K*_ as a function of their functional properties. Similarly, the connectivity kernel *w* describes the recurrent connectivity as a function of the neurons’ functional properties.

Second, we assume that the kernel *w* can be written in the form:

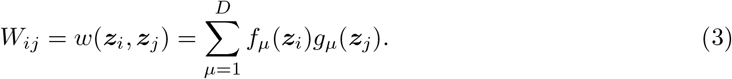

where *f*_1_, …, *f*_*D*_ and *g*_1_, …, *g*_*D*_ are arbitrary continuous functions over the similarity space ℳ (Fig. 2); and *D* ≪ *N* is a fixed number, not necessarily related to the dimension *d* of the similarity space. Motivated by results on low-rank RNNs, this assumption is rooted in the “neural manifold” perspective: in an RNN where the recurrent connectivity has rank *D*, the population activity lies in a *D*-dimensional neural manifold [58]. Models of low-rank networks can perform a wide range of computations, from the retrieval of patterns stored as fixed points (as in the Hopfield model) [64–66] to the approximation of arbitrary dynamical systems [38, 60]. We note that the matrix of recurrent connections in Eq.(3) can be written as the outer product of two *N* × *D* matrices *F* and *G*:

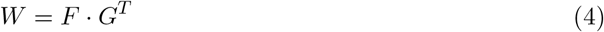

with entries *F*_*i,µ*_ = *f*_*µ*_(***z***_*i*_) and *G*_*j,µ*_ = *g*_*µ*_(***z***_*j*_). Therefore, we recognise the well-known structure of a connectivity with rank *D* (Fig. 2) [58].

**Fig. 2.**
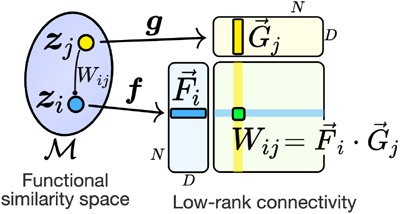
Linking circuit structure to low-rank connectivity. A postsynaptic neuron with location ***z***_*i*_ and a presynaptic neuron with location ***z***_*j*_ in the functional similarity space ℳ (left) are mapped by two functions, ***f***, ***g*** : ℳ → ℝ^*D*^, to two *D*-dimensional vectors 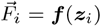 and 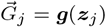, respectively. Each entry *W*_*ij*_ of the rank-*D* connectivity matrix (right) is given by the dot product 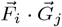.

Existing models of large low-rank networks assume that the rows {*F*_*i,µ*_}, {*G*_*i,µ*_} (*μ*= 1, …, *D*) of the matrices *F* and *G* in Eq.(4) are drawn independently across neurons *i* from a given distribution [58, 60, 61, 64–68]. In our framework, this distribution is (non-uniquely) parameterised by the density of neuronal locations ***z***_*i*_ in the similarity space, and by the choice of the functions ***f*** = (*f*_1_, …, *f*_*D*_) and ***g*** = (*g*_1_, …, *g*_*D*_). For instance, a Hopfield network with *P* binary patterns [64, 65] corresponds to locations ***z***_*i*_ uniformly distributed on the vertices {±1}^*P*^ of the *P*-dimensional hypercube and identity functions *f*_*µ*_(***z***_*i*_) = *g*_*µ*_(***z***_*i*_) = *z*_*i,µ*_ [69].

Thus, the assumption of Eq.(3) is a generalised formulation of the connectivity of large low-rank networks. This observation leads to our first result: models of large low-rank networks can generally be represented as circuit models, where each neuron is assigned a location in an underlying similarity space and the connectivity depends on the respective locations of the neurons as expressed by Eq.(3). If the locations ***z***_1_, …, ***z***_*N*_ of the neurons are drawn independently, the network dynamics of Eq.(2) then converges to the solution of a neural field equation in the large *N* limit [61, 70, 71], where the time-dependent membrane potential of each neuron *i* depends on its location ***z***_*i*_. (This convergence result actually holds for *all* networks with 1/*N* scaling of synaptic weights [72].) Therefore, we sometimes refer to a circuit model with many neurons as a “field model” in the rest of the paper.

For the ease of exposition – and without loss of generality (Methods 1) – we consider in the following that the rank *D* of the connectivity equals the dimensionality *K* of the input and that the functions (*f*_1_, …, *f*_*D*_) and 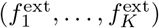 are the same. Our second theoretical result is that, if neuronal locations ***z***_*i*_ are sampled independently with probability distribution *ρ*(***z***), then, in the limit of a large number of neurons *N* → ∞, the potentials *h*_1_(*t*), …, *h*_*N*_ (*t*) can be written as

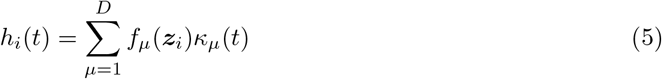

where the latent variables *κ*_1_(*t*), …, *κ*_*D*_(*t*) are the solution of

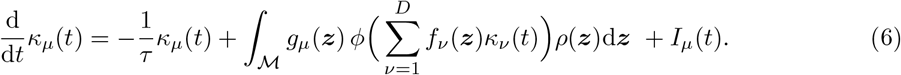

Equation (6) tells us three things. First, it is deterministic, implying that large networks of stochastic LNP neurons follow deterministic collective dynamics [61]. Second, the collective dynamics can be reduced to the nonlinear system of *D* ordinary differential equations given by Eq.(6), which means that the neural manifold is at most *D*-dimensional. Third, Eq.(6) expresses the dynamics of the latent variables *κ*_*µ*_(*t*), representing the coordinates of the population activity in the *D*-dimensional neural manifold, in terms of an integral over the *d*-dimensional similarity space ℳ. This provides an explicit mathematical link between the circuit structure, summarised by the locations of the neurons in the similarity space, and the flow of the collective dynamics on the neural manifold.

It is worth noticing that the low-rank connectivity matrix of Eq.(3) could be the result of Hebbian learning [64, 66]. Indeed, we can interpret each function *f*_*µ*_(***z***_*i*_) as the probability that neuron *i* with tuning properties ***z***_*i*_ forms, in the experimental condition *μ*, a dendritic spine, and *g*_*µ*_(***z***_*j*_) as the probability that neuron *j* forms a presynaptic bouton. The change of synaptic efficacy in condition *μ*will then be the product Δ*W*_*ij*_∝ *f*_*µ*_(***z***_*i*_)*g*_*µ*_(***z***_*j*_): across several experimental conditions, the low-rank connectivity of Eq.(3) will build up. In a memory-related brain area, the “experimental condition” could correspond to the presentation of a novel memory item; in early visual cortex, the experimental condition could correspond to elongated spatial edges observed during the critical period [73].

### Circuit structure and neural manifold can mismatch: examples

In this section, we highlight the insights provided by the above framework through a few examples. For the sake of simplicity, we assume, in these examples, that the network does not receive any external input and that the transfer function *ϕ* is a step function of the membrane potential *h*, i.e. *ϕ*(*h*) = *R* for *h*> 0 and zero otherwise. Qualitatively, results do not change for other choices of saturating transfer functions *ϕ* (Appendix B).

#### A one-dimensional circuit structure generating a two-dimensional neural manifold

In a first toy example, imagine that the functional properties of each neuron *i* are summarised by a single variable *z*_*i*_ ∈ ℝ, which is normally distributed across neurons (Fig. 3A1). This determines each neuron’s location in the one-dimensional similarity space ℳ = ℝ. If the interactions are given by *W*_*ij*_ ∝ *z*_*i*_*z*_*j*_, this corresponds to a Hopfield network with a single Gaussian pattern. In this case, the collective dynamics have a single stable non-trivial fixed point (up to a symmetry degeneracy) [60, 67]. We study a variant of this model where synaptic weights are given by:

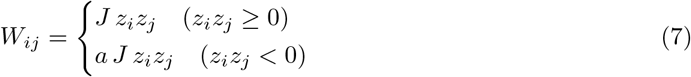

where *J*> 0 is the interaction strength, and negative weights (i.e. inhibitory synapses) are rescaled by a factor 0 < *a* < 1. The weights of Eq.(7) can be written in the form of Eq.(3), as: *W*_*ij*_ = *f*_1_(*z*_*i*_)*g*_1_(*z*_*j*_) + *f*_2_(*z*_*i*_)*g*_2_(*z*_*j*_) (Fig. 3A2), meaning that the connectivity has rank *D* = 2. The theory predicts two-dimensional collective dynamics, with three stable fixed points spread out in the plane and one saddle point at the origin (Fig. 3B; more details in Methods 2.1). This is confirmed by simulations of a SNN with *N* = 2000 neurons (obeying Eq.(1)) with weights given by Eq.(7). Thus, this model generates a neural manifold of dimension *D* = 2, whereas single-neuron responses depend on their abstract location in a (*d* = 1)-dimensional similarity space.

**Fig. 3.**
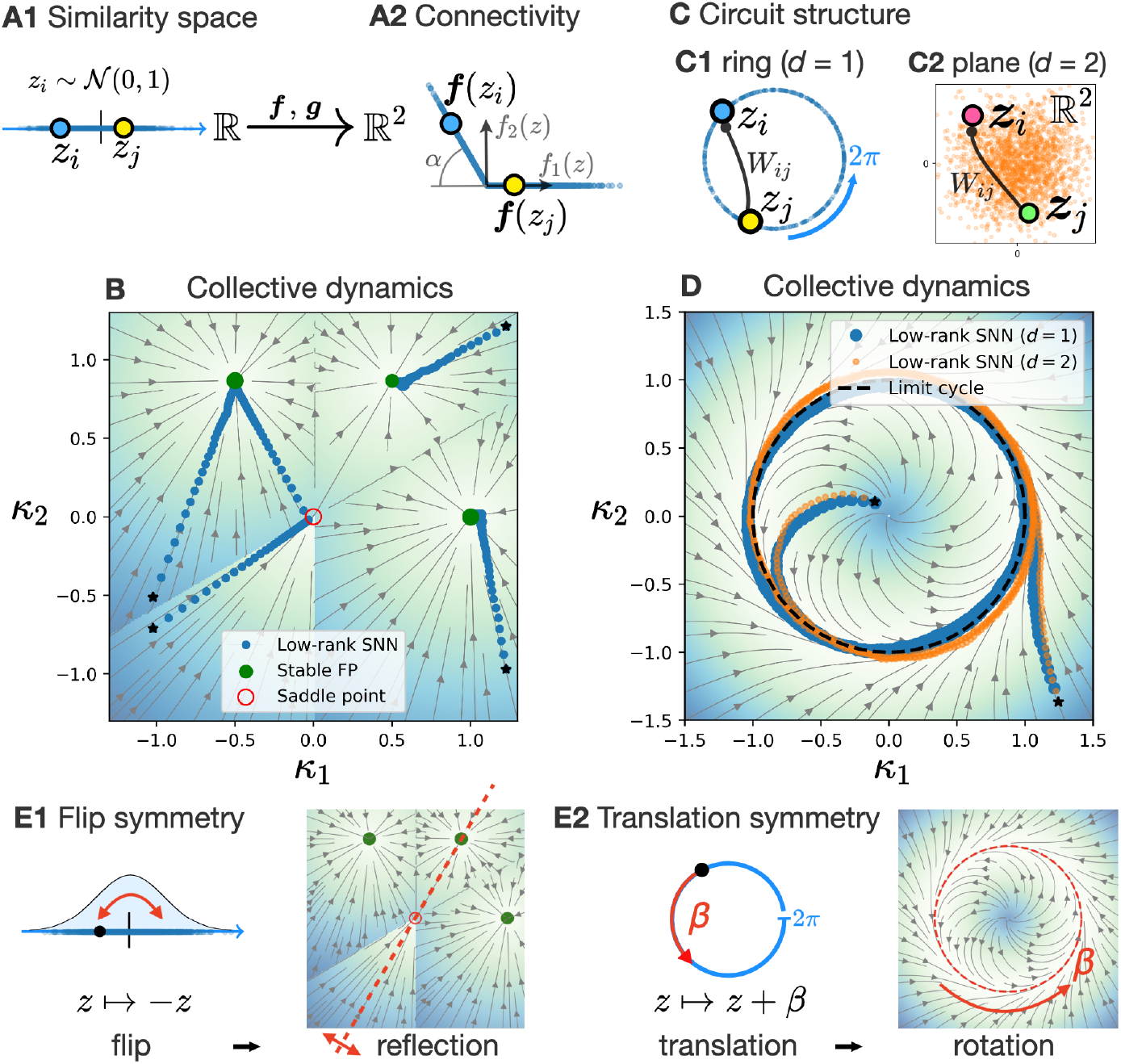
The dimension d of the functional similarity space and the dimension D of the neural manifold do not always match. **A-B**. One-dimensional circuit structure generating a two-dimensional neural manifold. **A1**. Neuronal locations are normally distributed over the real line: the dimension of the similarity space is *d* = 1. **A2**. The connectivity of Eq.(7) is given by 2-dimensional functions ***f*** and ***g*** = *J****f*** : the dimension of the neural manifold is *D* = 2. The angle of the kink is α = arccos *a*. **B**. Flow of the collective dynamics (Eq.(6)) of the circuit model in panel (A) (darker blue-shading indicates faster dynamics). The population trajectories of an SNN with *N* = 2000 neurons (blue dots), starting from 4 different initial conditions (black stars), follow the flow of the collective dynamics. The units of the axes are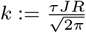. **C-D**. Two different circuit models generating identical limit-cyle collective dynamics. **C**. Schematic of the distribution of neuron locations: **C1** uniform over the ring (*d* = 1); **C2** standard Gaussian in the plane (*d* = 2). **D**. Same as panel (B), for the circuit models in panel (C), with the connectivity of Eqs.(8)-(9). The collective dynamics generate a globally attractive limit cycle. The units of the axes are 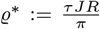 cos *δ*. **E**. *Microscopic* symmetries of the circuit structure induce *macroscopic* symmetries of the collective dynamics. **E1**. The microscopic symmetry w.r.t. a simultaneous flip of all neuron locations (flip symmetry) in panel (A) causes a reflection symmetry of the collective dynamics in panel (B). **E2**. The microscopic symmetry w.r.t. a simultaneous shift of all neurons along the ring (translation symmetry) in panel (C1) causes a rotation symmetry of the collective dynamics in panel (D). In panel (B), the SNN leaves the saddle point at the origin towards one of the attractors due to finite-size fluctuations and spike firing noise. The numerical parameters are: *τ JR* = 10, and *a* = 1/2 in (A-B), *δ* = *π*/10 in (C-D) (Eqs.(8)-(9)). The low-dimensional trajectory of the SNN is obtained from Eq.(1.8): 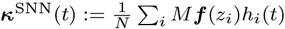

#### Two different circuit structures generating identical limit-cycle dynamics

We now turn to an example of limit-cycle dynamics, which is of practical relevance across different brain areas and species. For instance, in a task where a monkey has to grasp a lever and cycle with its hand, the activity in the motor cortex follows limit-cycle collective dynamics [30] that are likely generated by recurrent mechanisms [44]. In a different context, slow, internally-generated cycling dynamics have been recently observed in the mouse medial enthorinal cortex (MEC) [22] (Fig. 1D). We build here two models with different circuit structures that generate identical limit-cycle dynamics in a neural manifold of dimension *D* = 2. The first model is a *ring model*, in which neurons are uniformly assigned angular positions *z*_*i*_ ∈ [0, 2*π* [(Fig. 3C1) [16, 51]. The similarity space thus has dimension *d* = 1. We express the connectivity as:

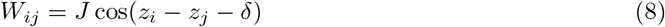

where *J*> 0 is the connectivity strength, and the angle *δ*∈] −*π, π*] is a phase shift parameter. For *δ* = 0, we get a standard ring attractor model with stationary bumps of activity [51], whereas for *δ* ≠ 0, the attractor states are travelling bumps [16]. Using the sum expansion of the cosine function [59], we see that the connectivity has rank *D* = 2, where the functions in Eq.(3) are given by *f*_1_(*z*) = cos *z, f*_2_(*z*) = sin *z* and *g*_*µ*_(*z*) = *Jf*_*µ*_(*z* + *δ*) (for *μ*= 1, 2).

In the second model, neurons have two-dimensional locations ***z***_*i*_ = (*z*_*i*,1_, *z*_*i*,2_) ∈ℝ^2^ normally distributed over the plane (Fig. 3C2); thus, the similarity space has dimension *d* = 2. As in the ring model above, the connections are maximal for an angle *δ* between two neurons:

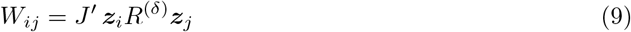

where *R*^(*δ*)^ is the 2 × 2 matrix of rotation by an angle *δ* in the plane, and *J*^*′*^ > 0 is the connectivity strength. The connectivity of Eq.(9) has also rank 2, since it is readily expressed as a two-dimensional dot product.

Remarkably, for 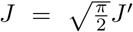 (i.e. up to a rescaling), the two models yield *identical* two-dimensional collective dynamics (Methods 2.1). The theory predicts that for any initial condition, the two latent variables that parameterise the population activity converge to a globally attractive limit cycle with angular speed *τ* ^−1^ tan(*δ*), which is confirmed by simulations of SNNs with *N* = 2000 neurons (Fig. 3D).

We highlight two aspects of this example. (i) Neither the dimension *D* of the neural manifold, nor the flow of the collective dynamics enable us to distinguish the ring model from the planar model. More generally, we found that a mismatch between the dimension of the similarity space and that of the neural manifold can originate from heterogeneity across the neuronal population: additional single-neuron properties, parameterising heterogeneity across neurons, can expand the dimensionality of the similarity space without affecting the collective dynamics. For instance, the above “planar” limit-cycle model can be interpreted as a ring model where each neuron is assigned an additional “radial” coordinate, independent of its angular location (Methods 2.2). Nevertheless, we present later, in the section “The footprint of circuit structure”, a method that allows us to extract circuit structure from neuronal activity. (ii) For both the “ring” and “planar” models, the connectivity is invariant under the simultaneous rotation of all neurons by any angle. In turn, the collective dynamics exhibits a rotation symmetry, visible in Fig. 3D. The general link between these symmetries is explained in the next section.

### Symmetries of circuit structure induce symmetries of collective dynamics

The general notion of a *symmetry* of a dynamical system refers to a transformation of the system that does not affect its dynamics. Our theory links circuit models and neural manifolds on the level of symmetries: symmetries of a circuit model, expressed as transformations of the neuronal locations in the similarity space, generate symmetries of the collective dynamics in the neural manifold. While this conceptual link is intriguing, its biological relevance is, at this stage, unclear. Nonetheless, in the following, we briefly present our main result and how it applies to the example models above.

Consider a simultaneous transformation of all neuronal locations, expressed as a mapping *S* : ℳ → ℳ over the similarity space that takes each neuron *i* from location ***z***_*i*_ to location *S****z***_*i*_. We call this transformation a *microscopic symmetry* if it commutes with the (microscopic) network dynamics of Eq.(2). Analogously, a *macroscopic symmetry* is a transformation of the latent variables (*κ*_1_, …, *κ*_*D*_), expressed as a mapping *T* : ℝ^*D*^→ ℝ^*D*^, that commutes with the collective dynamics of Eq.(6). In Methods 3 (and Appendix C), we show that each microscopic symmetry *S* of a circuit model induces a macroscopic symmetry *T* ^(*S*)^ of the associated collective dynamics in the *D*-dimensional neural manifold. For instance, the connectivity of the ring model, Eq.(8), is invariant under the translation of all neurons along the ring by any arbitrary (angular) distance β. This *microscopic* symmetry of the ring model is the origin of the evident *macroscopic* rotational symmetry of its collective dynamics (Fig. 3E2). The earlier example of a one-dimensional circuit model with connectivity given by Eq.(7) also has a microscopic symmetry, given by the “flip” *S* : *z* ↦ −*z* of each neuron’s position. The resulting macroscopic symmetry is the reflection in ℝ ^2^ w.r.t. the axis spanned by the vector 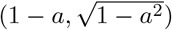 (Fig. 3E1).

### The footprint of circuit structure

We conclude from the previous sections that, despite the link provided by the symmetries, the functional similarity space and the neural manifold can have different dimensionality, and that several circuit models can generate the same collective dynamics.

Yet, even if the circuit structure provably can not be identified from the collective dynamics, can it be recovered by analysing neuronal activity? In this section, we propose an approach to this problem, that we apply on a model of context-dependent decision-making.

#### Single-neuron statistics reflect neuronal locations in the similarity space

Suppose that we have spiking data from an unknown circuit model network with *N* neurons. Using standard tools, we can extract at each time point the *N*-dimensional vector of the population activity, and trace out its trajectory on a *D*-dimensional neural manifold [27–29, 31–34]. On the other hand, in any circuit model with many neurons (i.e. a field model), the activity of each neuron *i* depends on its location in the functional similarity space. Thus, two neurons *i* and *j* at neighbouring locations ***z***_*i*_ ≈ ***z***_*j*_ will exhibit similar activity.

In order to extract similarity relations between neurons, we choose *T* different statistical measures that we apply to the spike train of each neuron: this yields an embedding of the neuron in *T* -dimensional space. Importantly, neighbourhood in the similarity space will lead to similar embeddings. The embeddings associated with several neurons will then reflect their relative positions in the similarity space. Consequently, if sufficiently many neurons are recorded, the *d*-dimensional similarity space will manifest itself as a point-cloud (with intrinsic dimension *d*) in the *T* -dimensional space of statistics. Thus, the above analysis should enable us to extract from *N*-dimensional spike data a *d*-dimensional similarity space, in which each neuron is assigned an empirical location.

In principle, neuronal embeddings could be generated using any well-chosen set of *T* statistical measures that summarise each neuron’s activity: for instance, it could be given by each neuron’s tuning to *T* different population-level factors; or by the “loadings” of the first *T* principal components (PCs) of population activity associated with each neuron [22] (as we illustrate in the next section). As a key property, these statistics should reflect functional similarities, so that neurons close to (far from) each other in the similarity space generate similar (different) outputs.

As an illustration, we first applied this method to extract the circuit structure of two different SNNs generating similar limit-cycle dynamics, with connectivity given as in the previous examples by Eqs.(8) and (9). We characterised each neuron’s spiking activity by its tuning to the two first PCs of the population activity, yielding a two-dimensional embedding of the whole similarity space (Methods 4 and Supp. Fig. S2).

### Retrieving circuit structure in models of context-dependent decision-making

Finally, we turned to a more relevant setup for systems neuroscience, in which model networks have to process external inputs to solve a given behavioural task. We applied our method for extracting the similarity space to simulated data obtained from two SNNs which solved the same task. This illustrates: (i) how our framework, linking circuit models and collective dynamics, can be used to build transparent models that solve a given task; and (ii) how different circuit models can be characterised by extracting their similarity space from the neuronal activity.

The chosen task is a simplified instance of context-dependent decision-making [34]. During a given trial, the network receives a sensory input consisting of two independent signals, named ‘color’ and ‘motion’, which can both be either positive or negative; as well as a binary context input which indicates which of the ‘color’ or ‘motion’ signal is relevant (Fig. 4A). The task consists in indicating, in the ‘color’ context, whether the ‘color’ signal is positive or negative at trial onset, and similarly for the ‘motion’ context. The ‘choice’ of the network is given by a weighted readout of its low-passed spiking activity.

**Fig. 4.**
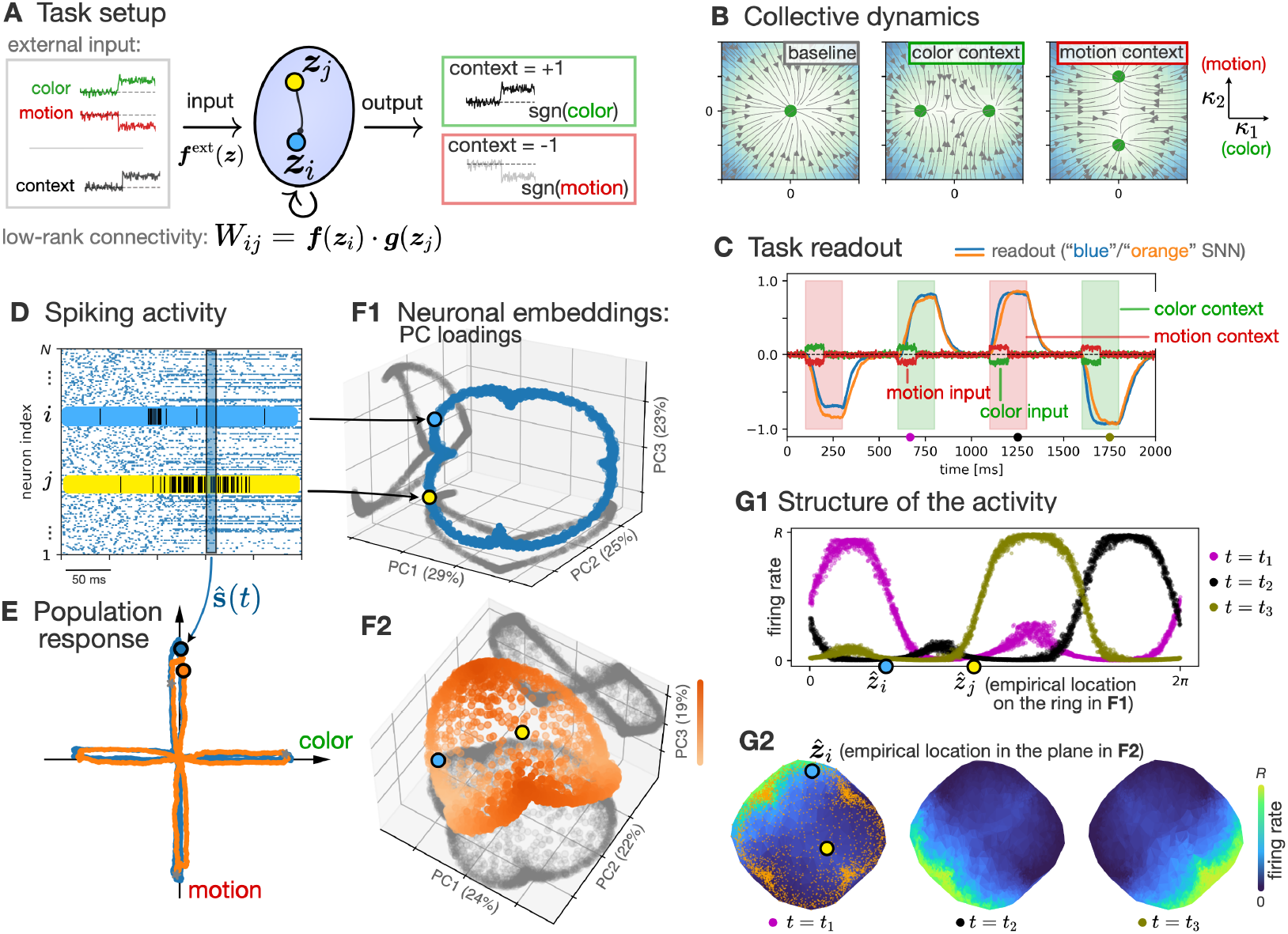
Extracting the circuit structure of two SNNs performing the same task. **A**. During each trial, networks receive an external input consisting of two sensory features (named ‘color’ and ‘motion’), each fluctuating around a positive or negative mean value, and a binary context feature. The task is to indicate the sign of the sensory feature that is specified by the context (‘color’ for the context being +1, and ‘motion’ for −1). Input, recurrent, and output synaptic weights are determined by the neurons’ locations {***z***_*i*_} in an underlying similarity space. **B**. The task is solved by two-dimensional collective dynamics, where the context input generates attractors (green dots) along an axis corresponding to the relevant sensory input feature. This is implemented in two SNNs, called “blue” and “orange”, of *N* = 3000 neurons each **C**. Time-course of the sensory inputs (green and red traces for the ‘color’ and ‘motion’ features) and normalised readout of the two SNNs (blue and orange traces) during 4 trials. The context input takes value +1 (respectively − 1) in the ‘color’ (resp. ‘motion’) context, indicated by green (resp. red) shadings, and 0 in between trials. Both networks correctly solve the task. **D**. Spike trains of 150 example neurons from the “blue” SNN. Spike trains of example neurons *i* and *j* are highlighted. **E**. Low-dimensional trajectory of the two SNNs (blue and orange traces) during 4 trials, obtained by projecting the *N*-dimensional low-passed spiking activity **ŝ**(*t*) onto ‘color’- and ‘motion’-relevant axes. **F**. The spiking activity of each neuron is characterised by the corresponding loadings on the three first principal components (PCs) of the population activity, which determines an embedding of the neuron in three dimensions. **F1**. For the “blue” SNN, the point-cloud of the embeddings of all the neurons in the population forms a ring, revealing the similarity space of the circuit model. **F2**. In contrast, for the “orange” SNN, the set of all three-dimensional embeddings lies on a curved, two-dimensional surface. Points are coloured according to their height (loading on PC3). **G1**. Each neuron *i* in the “blue” SNN is assigned an empirical location 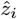 according to its position on the ring-like point-cloud in panel F1. Coloured curves represent the neurons’ momentary firing rates as a function of their empirical locations, at three different time points *t*_1,2,3_ (coloured dots on the x-axis of panel C). **G2**. Each neuron *i* in the “orange” SNN is assigned a two-dimensional empirical location 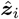 according to its position on the two-dimensional point-cloud in panel F2. Heatmaps represent the neurons’ firing rates as a function of their empirical locations, at the same three time points *t*_1,2,3_ (from left to right). Orange dots on the left-most heatmap represent all empirical locations inferred from panel F2. In panels F1-2, the percentage of variance of the population activity explained by each PC is indicated on the corresponding axis. Grey point-clouds show projections of the points onto the horizontal and a vertical plane. Details about implementation in Methods 4.2-4.3.

We designed two SNNs with different circuit structures, composed of *N* = 3000 neurons each, that we will call the “blue” and the “orange” SNN, which both solve the task through similar two-dimensional collective dynamics [58] (Methods 4). For both models, population activity decays towards a baseline state between two trials, while in the ‘color’ context, symmetric attractive fixed points appear along a ‘color’-relevant axis, and similarly in the ‘motion’ context (Fig. 4B). We aim to identify the circuit structure of each of the two SNNs based on their activity.

#### Extracting circuit structure from neuronal activity

The two-dimensional trajectory corresponding to the population activity of each SNN across four trials was obtained by projecting its low-passed, *N*-dimensional spiking activity onto ‘color’ and ‘motion’-relevant axes [34]. Upon receiving the same sensory and context inputs, the low-dimensional trajectories and the choice readouts of the two SNNs were similar (Fig. 4C,E).

A 3-dimensional embedding of each neuron was then obtained by measuring, for each neuron, the associated loadings onto the first three principal components (PCs) of the population activity (Fig. 4F). For the “blue” SNN, the point-cloud of the embeddings of all the neurons in the population forms a ring; whereas for the “orange” SNN, it lies on a curved, 2-dimensional surface. For both networks, the similarity space inferred from statistical features reflects the underlying circuit structure: the “blue” SNN corresponds to a ring model, and the “orange” SNN to a model with a 2-dimensional similarity space (Methods 4). Neurons were then assigned empirical locations based on their relative positions in the inferred similarity space. A representation of the neuronal activity as a function of the empirical neuronal locations highlights its underlying spatial structure, which differs between the two networks (Fig. 4G).

#### Extracting a task-irrelevant circuit structure component

In the previous example, the difference between the circuit model underlying each of the two “blue” and “orange” SNNs could have been detected by inspecting the tuning of each neuron to the sensory input features. Suppose that we assign to each neuron a two-dimensional “tuning vector”, that characterises its response to the two input features ‘color’ and ‘motion’. In both models, individual neurons are tuned to mixtures of these features (“mixed selectivity” [14]). Yet for the “blue” network, the tuning of each neuron to these two features is correlated, so that the corresponding set of two-dimensional tuning vectors lies on a circle; which is not the case for the “orange” network.

Because our approach is task-agnostic, a circuit structure component that is irrelevant to the task at hand can also be extracted. As an illustration, we considered a third “pink” SNN, where the ring circuit structure of the “blue” SNN described above was modified by the addition of a task-irrelevant, Hebbian-like connectivity component (Fig. 5A). To do this, a task-irrelevant feature *ξ*_*i*_ = ±1 was assigned to each neuron *i* (chosen independently across neurons with equal probability of being +1 or − 1); and a term proportional to *ξ*_*i*_*ξ*_*j*_ was added to the synaptic weight between pairs of neurons *i* and *j* (Methods 4). The functional similarity space of this new “pink” model now consists of two rings: each neuron *i* is jointly characterised by an angular position *z*_*i*_, and by the binary value of the additional feature *ξ*_*i*_. Since this additional feature is irrelevant to the task, it cannot be detected through the tuning of the neurons to the task variables (context, color or motion). If both SNNs receive identical external inputs, their respective population activity follow similar low-dimensional trajectories along task-relevant axes (Fig. 5B).

**Fig. 5.**
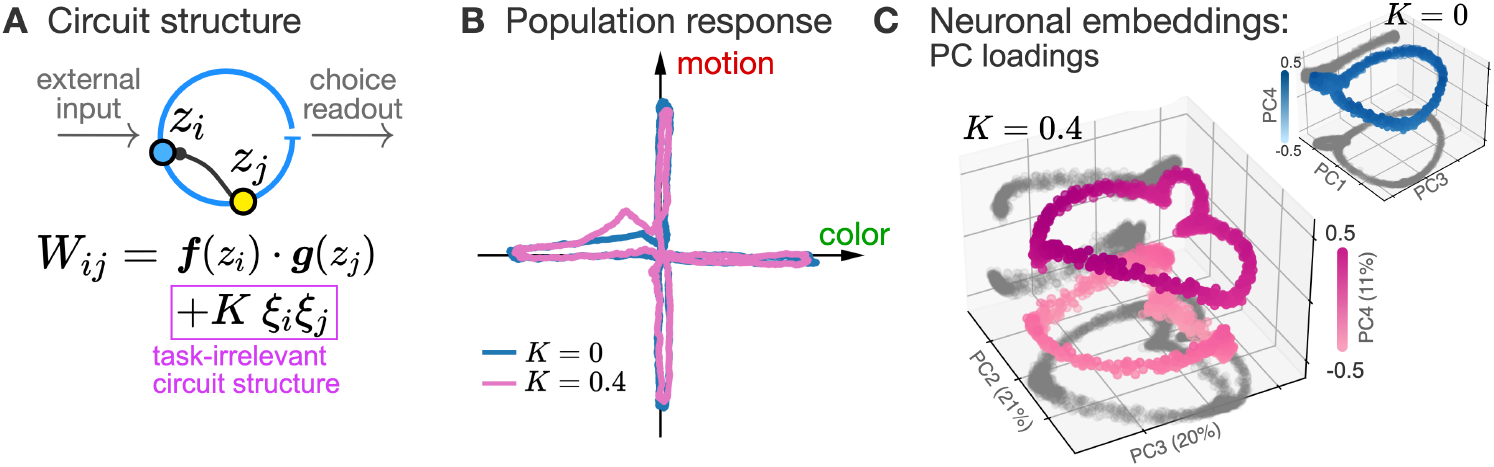
Extracting a task-irrelevant circuit structure component from population activity. **A**. Two SNNs (of *N* = 3000 neurons each) solve the same task as depicted in Fig. 4A. The first SNN corresponds to a ring model (the “blue” SNN of Fig. 4). In the second “pink” SNN, in addition to its location *z*_*i*_ on the ring, each neuron is assigned a task-irrelevant binary feature *ξ*_*i*_ = *±*1, distributed independently across neurons; and a task-irrelevant component *Kξ*_*i*_*ξ*_*j*_, of strength *K* = 0.4 > 0, is added to the synaptic weights between pairs of neurons *i, j*. **B**. Low-dimensional trajectories of the two SNNs (blue solid line for the original “blue” SNN, for which *K* = 0; and pink solid line for the “pink” SNN with *K* = 0.4) across 4 trials during which they received the same external input, obtained by projecting the *N*-dimensional spiking activity on two task-relevant axes (as in Fig. 4D-E). **C**. 3-dimensional embeddings of the neurons in the “pink” (*K* = 0.4) and the “blue” SNN (*K* = 0; small inset). For the “pink” SNN, the three-dimensional point-cloud forms two disjoint rings, revealing the similarity space of the circuit model. Points on the lower or upper ring correspond to neurons with a task-irrelevant feature *ξ*_*i*_ =− 1 (resp., *ξ*_*i*_ = +1). The embedding of each neuron is given by the corresponding loadings on the PCs 2,3 and 4 of the population activity (PCs 1,3,4 for the “blue” SNN). Points are coloured according to their projection onto the vertical axis. Plotting conventions are as in Fig. 4F. Inset: the single-ring circuit structure of the original “blue” SNN is recovered for *K* = 0 (similar to Fig. 4F1).

Low-dimensional embeddings of the neurons in both networks were obtained as in Fig. 4 by measuring, for each neuron, the associated loadings onto the first PCs of the population activity. The resulting point-clouds reflect the underlying circuit structure: for the “pink” SNN, the pointcloud forms two disjoint rings, revealing the topology of the similarity space (Fig. 5C); whereas one recovers the original single-ring structure for the “blue” SNN.

## Discussion

To address the relationship between the functional similarity space, assumed to reflect the structure of cortical circuits, and the neural manifold, assumed to characterise computations via the collective dynamics of population activity, we formulated three specific questions in the Introduction. The first question focused on the mathematical link between the structured connectivity of a circuit model and the flow of its collective dynamics in the neural manifold. To answer this question, we proposed a modelling framework that explicitly provides this link. Our framework relies on the key idea that, in the limit of large networks, both low-rank RNNs and classical circuit models (where the connectivity is structured by the functional properties of the neurons) can be interpreted in the general framework of neural field models. In passing, we answered the question of whether “low-rank RNNs involve mechanisms similar to classical circuit models or implement truly novel solutions” [3]. The second question, addressing whether a unique relationship exists between the functional similarity space and the neural manifold, must be answered negatively: different circuit structures can yield equivalent collective dynamics. On the other hand, we proved that symmetries of the circuit structure give rise to symmetries of the collective dynamics, a result that highlights the importance of symmetries for models of computation in recurrent networks.

The third question explored the possibility of extracting knowledge about the circuit structure from recordings of neuronal activity. As an answer to this question, we proposed a method for extracting the topology of the similarity space of a circuit model from the neuronal activity, and applied it on simulated data. Our method allowed us to distinguish between different circuit structures giving rise to the same collective dynamics and solving the same behavioural task. In contrast with the analysis of neuronal tuning or selectivity, this approach is task-agnostic, which allowed us to detect a circuit structure irrelevant to a given behavioural task. In conjunction with task- and behaviour-informed methods [32], the unsupervised approach we propose for characterising functional similarities could enable the discovery of intricate features that modulate neuronal responses in higher cortical areas, and could also be applied in the absence of any task on recordings of freely behaving animals [74, 75].

While our modelling framework provides a link between circuit structure and low-dimensional neuronal activity, it does not incorporate important features of more realistic models of biological neural networks. First, in our model, recurrent connectivity and neuronal properties are constructed according to a homogeneous structural principle (that of field models), whereas cortical neuronal responses exhibit a large degree of trial-to-trial variability [76, 77], which could be explained by chaotic or chaos-like activity due to randomness in the connectivity [58, 66, 78]. The generalisation of our theory to the case of mixed structured and random connectivity, following [58], is left for future work. Second, the neurons in our model are linear-nonlinear-Poisson neurons, which do not account for neuronal refractoriness. Our theory can be partly extended to models of renewal neurons (such as the Spike Response Model or Integrate-and-Fire models [79, 80]). Because of the reset, neuronal dynamics is no longer memory-free and requires an extended formalism [79]. Yet, in the regime of asynchronous firing, each neuron’s firing rate can be approximated by a non-linear function of its synaptic input [81], so that our formalism can be applied to the description of the fixed points of the asynchronous population activity. Third, cortical networks exhibit a balance between excitation and inhibition [82, 83]. This has been taken into account in models of balanced networks [84–87], which can include a spatial structure [88, 89]. Whether our formalism can be extended to derive a low-dimensional description of spatially structured balanced networks remains an open problem.

Finally, we did not apply our method to extract the functional similarity space from experimental data. Yet, in ref. [22], an approach similar to that proposed in this paper was used on recordings of the mouse medial enthorinal cortex: single-neurons were characterised by a few statistics (the PC loadings) extracted from the population activity, and assigned a location on a ring according to these statistics (similar to Supp. Fig. S2); this revealed a bump of neuronal activity slowly cycling over this virtual ring. The question of whether such a low-dimensional population structure can be generally observed in experimental recordings remains to be tested. A positive answer would mean that, even in a population where neurons exhibit mixed selectivity and seemingly heterogeneous activity, neuronal responses can be parameterised by only a few variables, making field models well-suited for describing collective neuronal activity. More broadly, the link between neural manifolds and circuit (or field) models established in this work could potentiate the use of large-scale recordings to uncover circuit structures underlying population-level computation in large networks of neurons.

## Methods

### 1 A unifying framework: field model of low-rank networks

#### Network model

We turn here to a more precise mathematical formulation. We consider a model of Recurrent Neural Network (RNN) composed of *N* linear-nonlinear-Poisson neurons [62]. The spike train S_*i*_(*t*) of each neuron *i* = 1, …, *N* is generated by an inhomogeneous Poisson process with intensity *r*_*i*_(*t*) = *ϕ*(*h*_*i*_(*t*)), where *h*_*i*_(*t*) represents the neuron’s membrane potential, and *ϕ* : ℝ → ℝ ^+^ is a non-linear bounded voltage-to-rate function. The potential *h*_*i*_(*t*) is a leaky integrator of the synaptic input; which, for the ease of exposition, we first consider as coming only from recurrent connections. The dynamics of the potential thus follow:

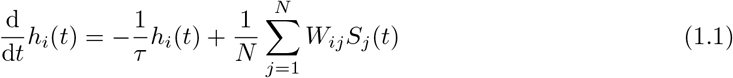

with given initial condition *h*_*i*_(0). Here, *τ*> 0 is the membrane time constant; and the synaptic weights, scaled as 1/*N*, are given by a matrix of rank *D* ∈ ℕ fixed:

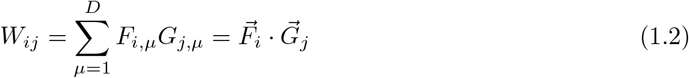

where the dot denotes the dot product in ℝ ^*D*^. Each entry *W*_*ij*_ of the low-rank connectivity matrix corresponds to the dot product of two *D*-dimensional “connectivity vectors”, 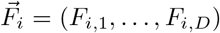 and 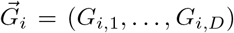, associated with neurons *i* and *j* (Fig. 2). As discussed in the Results section, Eq.(1.2) is a general formulation of the constraint of a low-rank connectivity; specific models of low-rank networks are essentially built by specifying the distribution of these vectors across the population [58, 60, 61, 64–68]. In what follows, we introduce a general model for the distribution of the connectivity vectors. The resulting class of RNNs encompasses most known models of mean-field low-rank networks. Moreover, for fixed rank *D*, as *N*→ ∞, it corresponds to a field model over a similarity space whose dimension *d* is not necessarily related to the rank *D* of the connectivity, i.e. to the dimension of the neural manifold.

#### A general model of low-rank connectivity

As we are interested in the limit of large *N*, let’s start from the simple observation that, if one wants to add a neuron labelled *i*^∗^ to the network, one has to specify its connectivity, through the associated vectors 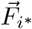 and 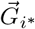, as well as its initial condition *h*_*i*∗_ (0). It is commonly assumed that these features are independent across neurons: intuitively, it means one can add a neuron *i*^∗^ to the network without knowing the 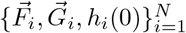 associated with all other neurons. However, for a given neuron *i*, the components of 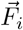 and 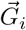 might be correlated. For the new neuron *i*^∗^, we thus model these features as being determined by some *functional properties* of the neuron, sampled from a given distribution.

Formally, we represent these functional properties by the neuron’s *location* ***z***_*i*∗_, drawn at random according to a probability distribution *ρ*(d***z***) on a smooth manifold ℳ of intrinsic dimension *d*. Given the location ***z***_*i*_ of any neuron *i*, the associated connectivity vectors 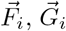 are determined by mappings ***f***, ***g*** : ℳ → ℝ ^*D*^:

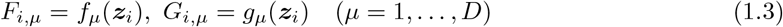

with *f*_*µ*_, *g*_*µ*_ the *μ*-th components of ***f***, ***g***. This is illustrated on Fig. 2. Analogously, the initial condition is determined by a field *h*_0_ : ℳ → ℝ, according to *h*_*i*_(0) = *h*_0_(***z***_*i*_).

Eqs.(1.2),(1.3) define the connectivity kernel *w*(***z, z***^*′*^) = ***f*** (***z***) · ***g***(***z***^*′*^), such that the recurrent connectivity is given by *W*_*ij*_ = *w*(***z***_*i*_, ***z***_*j*_). The manifold ℳ thus represents the functional similarity space, in which the neurons’ locations determine the connectivity. The distribution of the neuronal locations, described by *ρ*(d***z***), defines an inner product between functions *φ, ψ* over ℳ:

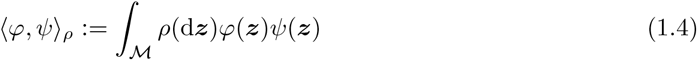

In the following, we assume that the connectivity functions *f*_*µ*_, *g*_*µ*_ (*μ*= 1, …, *D*), as well as the initial condition *h*_0_, are continuous and belong to the vector space of *ρ*-square-integrable functions over ℳ. This last condition defines a Hilbert space ℋ equipped with the *ρ*-inner product ⟨·, ·⟩_*ρ*_.

#### Corresponding field model

As *N* → ∞, the RNN defined by Eqs. (1.1,1.2,1.3) is described by a field model: the membrane potential at time *t* of a neuron at location ***z*** is given by the neural field v_*t*_(***z***), which obeys the *field equation*

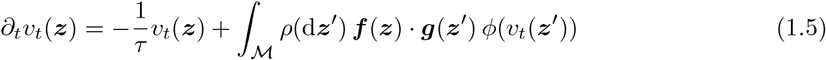

with initial condition v_0_(***z***) = *h*_0_(***z***). It is the continuum version of Eq.(1.1) [63, 70], where the Poisson noise from the spikes vanishes [61, 71]; i.e., S_*i*_(*t*) was replaced by the expected instantaneous rate *ϕ*(*h*_*i*_(*t*)). Under suitable regularity and boundedness assumptions concerning the transfer function *ϕ*,the convergence as *N* → ∞ of each membrane potential *h*_*i*_(*t*) to v_*t*_(***z***_*i*_) is guaranteed, with rate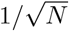. It follows that neurons become pairwise independent [71, 90].

#### Low-dimensional dynamics

Analogously to a rank-*D* RNN [60], the dynamics of the neural field can be written as a *D*-dimensional system [59], as presented here in a simple fashion. The field *v*_*t*_, at any time *t*, is an element of the Hilbert space ℋ (provided that *h*_0_, {*f*_*µ*_}_*µ*_ ⊂ ℋ). The field equation (1.5) can thus be written as an ODE in ℋ: [91]

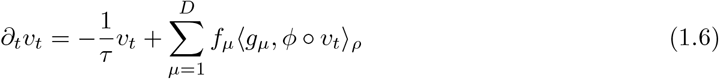

where (∘) denotes the composition of functions, and ⟨·, ·⟩_*ρ*_ the *ρ*-inner product (Eq.(1.4)). The field v_*t*_ undergoes an exponential decay, except in the linear subspace *F* := span{*f*_*µ*_} ⊂ ℋ. It can thus be written as a linear combination of the basis functions {*f*_*µ*_}, whose coefficients *κ*_*µ*_(*t*) are the “latent variables”:

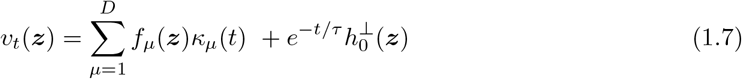

where 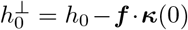 is the projection of the initial condition *h*_0_ onto the orthogonal complement of ℱ inℋ. By projection of Eq.(1.7) onto each basis function *f*_*µ*_, the latent variables can be written in vector form as:

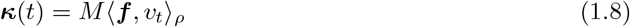

where we introduced the inverse correlation matrix ℳ, defined by (ℳ ^−1^)_*µν*_ = ⟨*f*_*µ*_, *f*_*ν*_⟩_*ρ*_ (assuming without loss of generality that the functions {*f*_*µ*_} are linearly independent in ℋ). Eqs. (1.6-1.7-1.8) yield *D*-dimensional collective dynamics for ***κ***, which can be written compactly:

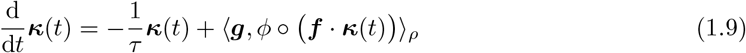

under the assumption that 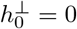. This equation for the collective dynamics (and its more general expression, Eq.(A.13) in the Appendix) is the central ingredient of our theory. The derivation is given in Appendix A.

#### Neural manifold and attractor states

Eq.(1.7) implies that the firing rate of each neuron *i*, given by *r*_*i*_(*t*) = *ϕ*(*h*_*i*_(*t*)) = *ϕ*(v_*t*_(***z***_*i*_)), is parametrised by the latent variables ***κ***; and so is the *population activity* **r**(*t*), which is the joint vector of all *N* instantaneous firing rates *r*_1_(*t*), …, *r*_*N*_ (*t*). Besides, **r**(*t*) exponentially decays towards the *D*-dimensional neural manifold

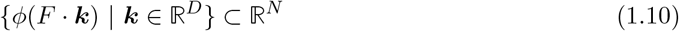

generated by the set of all possible latent variables; where *F* ∈ ℝ ^*N×D*^ denotes the matrix of connectivity vectors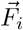 given by Eq.(1.3), and *ϕ* is applied element-wise. In the absence of external input, the latent variables ***κ***(*t*) will converge to the set of attractor states of the collective dynamics of Eq.(1.9), which specifically depends on the considered model. In the Results, we give concrete examples, in which attractor states are either discrete fixed points or limit cycles.

#### Generalisations: external input and arbitrary connectivity kernel

This derivation can be readily extended to the case of a network receiving a low-dimensional external input. We assume that the external input is composed of *K* features *I*_*ν*_(*t*) (*ν* = 1, …, *K*), and that the input weight to a neuron *i* from the input feature *I*_*ν*_ is determined by a corresponding function 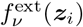 of the neuron’s location ***z***_*i*_. The external input it receives can thus be written:

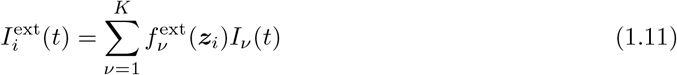

This simply results in a drive to the collective dynamics of Eq.(1.9), while expanding their dimensionality – at most up to *D* + *K*. The corresponding general expression of the collective dynamics is given by Eq.(A.13) in Appendix A.2.

The derivation of the collective dynamics can also be generalised to the case of field models with arbitrary square-integrable interaction kernel over ℳ^2^. In this case, the collective dynamics are not guaranteed to be finite-dimensional; but the field can always be well approximated by a low-rank model by a truncation of the dimension [59]. Both generalisations are given in Appendix A.

## 2 Similarity space and neural manifold can mismatch

### 2.1 Examples

#### A one-dimensional circuit structure generates two-dimensional collective dynamics

In our first example, neuronal locations are normally distributed over the real line; i.e. ℳ = ℝ and *z*_*i*_ ∼ *ρ* = *N*(0, 1) i.i.d. for *i* = 1, …, *N* (Fig. 3A1). The interactions are given by Eq.(7):

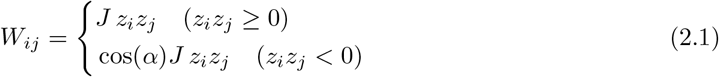

with 0 < α < *π*/2 (and *a* = cos α in Eq.(7)). This can be written as the 2-dimensional dot product *W*_*ij*_ = ***f*** (*z*_*i*_) · ***g***(*z*_*j*_), with:

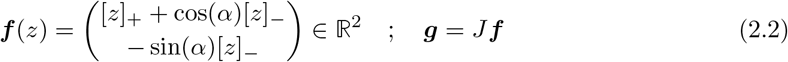

where [*z*]_*±*_ = *z* · 𝟙_{sgn(*z*)=*±*1}_ denotes the positive or negative part of *z*. This is illustrated on Fig. 3A2. The corresponding collective dynamics resulting from Eq.(1.9) and displayed in Fig. 3B are derived in Appendix B.1.

#### Two different circuit structures generate identical limit-cycle dynamics

We consider two circuit models, with connectivity given by Eqs.(8)-(9), which we recall here for convenience. The first model is a ring model, with angular locations *z*_*i*_ uniformly distributed on the ring [0, 2*π*[, and rank-2 connectivity given by:

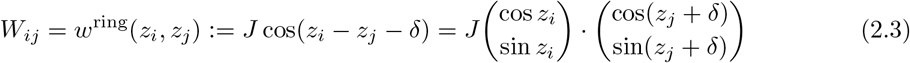

with *δ* an arbitrary phase shift. In the second “planar” model, neurons have two-dimensional locations ***z***_*i*_ = (*z*_*i*,1_, *z*_*i*,2_) ∈ ℝ ^2^ normally distributed over the plane; and synaptic weights are given by:

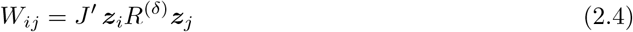

where *R*^(*δ*)^ is the 2 × 2 matrix of rotation by an angle *δ* in the plane; a nd *J*^*′*^ > 0.

Remarkably, for a step transfer function *ϕ*(x) = *R*· 𝟙_{*x>*0}_, and 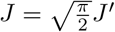the two models obey *identical* collective dynamics. For the polar coordinates (*ϱ*(*t*), *θ*(*t*)) of the latent variable ***κ***(*t*) ∈ ℝ ^2^, Eq.(1.9) yields

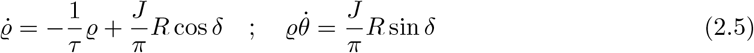

for *both* models (derivation in Appendix B.2).

For any initial condition, the latent variables converge to a globally attractive limit cycle with constant angular speed which only depends on *τ* and *δ* (given by 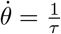 tan *δ*). This last point generalises to arbitrary transfer function *ϕ* and any rotationally invariant distribution of the neuronal locations (Appendix B.2).

### 2.2 General interpretation

#### Heterogeneity as a source of dimensionality mismatch

We observed that the dimensionality *d* of the similarity space can be expanded by additional single-neuron properties, that parameterise heterogeneity across the population. As a concrete example, consider the “planar” limit-cycle model above with a two-dimensional similarity space (Eq.(2.4)). In this example, neuronal locations are normally distributed in ℝ ^2^. Thus, the location ***z***_*i*_ of any neuron *i* can be written in polar coordinates, with an angle*φ*_*i*_ (distributed uniformly over the ring) and a radius *ξ*_*i*_ (distributed following the χ_2_-distribution, independent from*φ*_*i*_). The connectivity of Eq.(2.4) then reads *W*_*ij*_ = *ξ*_*i*_*ξ*_*j*_*w*^ring^(*φ*_*i*_,*φ*_*j*_), where *w*^ring^ denotes the connectivity kernel of the ring model (Eq.(2.3)). We now interpret the radius *ξ*_*i*_ as a single-neuron property that parameterises an heterogeneity across the population, affecting the connectivity of the original ring model (for which *ξ*_*i*_ = 1 for all neurons). Yet, this heterogeneity does not affect the collective dynamics of the model (Appendix B.2). Building on this observation, we found that additional single-neuron properties that parameterise heterogeneity across the population can be incorporated into any circuit model, thus extending the dimensionality *d* of the similarity space, without affecting the collective dynamics. Simple examples include the multiplication of all outgoing weights of each neuron by a random factor, that describes variations of synapse strength; and heterogeneous maximum firing rates across neurons (Appendix D).

#### Several universal approximators

The mismatch between the circuit structure and the collective dynamics can also be understood from another perspective: several circuit models with different similarity spaces can approximate the same collective dynamics. We identified two types of circuit models that are “universal approximators”. On the one hand, a Gaussian mixture low-rank network of rank *D* can approximate arbitrary collective dynamics in a *D*-dimensional neural manifold [60]. This class of models has a functional similarity space of dimension *d* = 2*D* (Appendix E.1). On the other hand, field models with a one-dimensional similarity space can in principle also approximate arbitrary collective dynamics, if the functions *f*_*µ*_, *g*_*µ*_ in Eq.(3) can be chosen arbitrarily with no requirement on their “smoothness” – analogous to the universal approximation property of feed-forward neural networks (Appendix E.2).

## 3 Symmetries

A *microscopic symmetry* of a circuit model can be informally defined as follows. Consider a transformation of the neuronal locations, expressed as a bijection *S* : ℳ → ℳ over the similarity space that takes each neuron *i* from location ***z***_*i*_ to location S***z***_*i*_. There are two ways in which this mapping can be combined with the microscopic network dynamics of Eq.(2). Given the set of all membrane potentials at initial time, one can first let the network evolve up to time *t* and then “move” each neuron *i* from location ***z***_*i*_ to S***z***_*i*_. Alternatively, one can first “move” the neurons and then let the network evolve up to time *t*. A microscopic symmetry of the model is a transformation *S* of the neuronal locations for which these two orders of composition are equivalent (Fig. S1).

Analogously, a *macroscopic symmetry* is a transformation of the latent variables (*κ*_1_, …, *κ*_*D*_) that commutes with the collective dynamics of Eq.(6).

### A symmetry is a mapping that commutes with the flow

A more precise formulation relies on the concept of *flow* of the dynamics. Consider that the membrane potential of each neuron is given, at initial time *t* = 0, by a function *v*_0_ of its location; i.e., *h*_*i*_(0) = *v*_0_(***z***_*i*_). The *flow* of the dynamics transforms this initial condition into the function *v*_*t*_ that describes the membrane potentials at time *t* (i.e. *h*_*i*_(*t*) = *v*_*t*_(***z***_*i*_)). In other words, the flow is a mapping Φ_*t*_ such that v_*t*_ =Φ_*t*_[*v*_0_] (Fig. S1A, left panels). On the other hand, if one “moves” each neuron from location ***z***_*i*_ to location *S****z***_*i*_, its membrane potential is now given by the value of the field at its “new” position; i.e. one now has *h*_*i*_(0) = *v*_0_(*S****z***_*i*_). In other words, “moving” the neurons is equivalent to a composition of the field with *S* (Fig. S1A, top panels). As described above, the bijection *S* is a *microscopic symmetry* of the model if “moving” the neurons before or after the time evolution is equivalent; i.e.,Φ_*t*_[*v*_0_ ° *S*] = Φ_*t*_[*v*_0_] ° *S*.

Likewise, a *macroscopic* symmetry of the model is a transformation *T* : ℝ ^*D*^ → ℝ ^*D*^ of the latent variables (*κ*_1_, …, *κ*_*D*_) that commutes with the flow of the collective dynamics of Eq.(6) (Fig. S1B).

### From microscopic to macroscopic symmetries

We show that any microscopic symmetry *S* of a field model generates a macroscopic symmetry *T* ^(*S*)^ of its collective dynamics. For the latter, we give an explicit expression under a reasonable additional assumption. More precisely (Theorem 1 in Appendix C):

1. Let *S* be (i) a microscopic symmetry of the field model, and (ii) under which the *D*-dimensional space 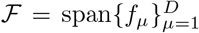 containing the recurrent input is invariant. This is expressed by the condition: *f*_*ν*_ ° *S* ∈ *F*, for all *ν* = 1, …, *D*. Then, a symmetry of the collective dynamics is given by the linear mapping *T* ^(*S*)^, defined by the *D* × *D* matrix

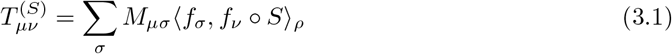

with ℳ the inverse correlation matrix, defined by (*M* ^−1^)_*µν*_ = ⟨*f*_*µ*_, *f*_*ν ρ*_⟩.
2. If *S* is a bijection for which both the connectivity kernel *w* and the distribution of the locations *ρ* are invariant, i.e. *w*(*Sz, Sz*^*′*^) = *w*(*z, z*^*′*^) and *ρ* ∘ *S*(*dz*) = *ρ*(*dz*), then both requirements (i) and (ii) of point 1. above are satisfied.

*A fortiori*, any symmetry *T* ^(*S*)^ of the collective dynamics will be a symmetry of the set of attractor states of the population activity.

**Fig. S1.**
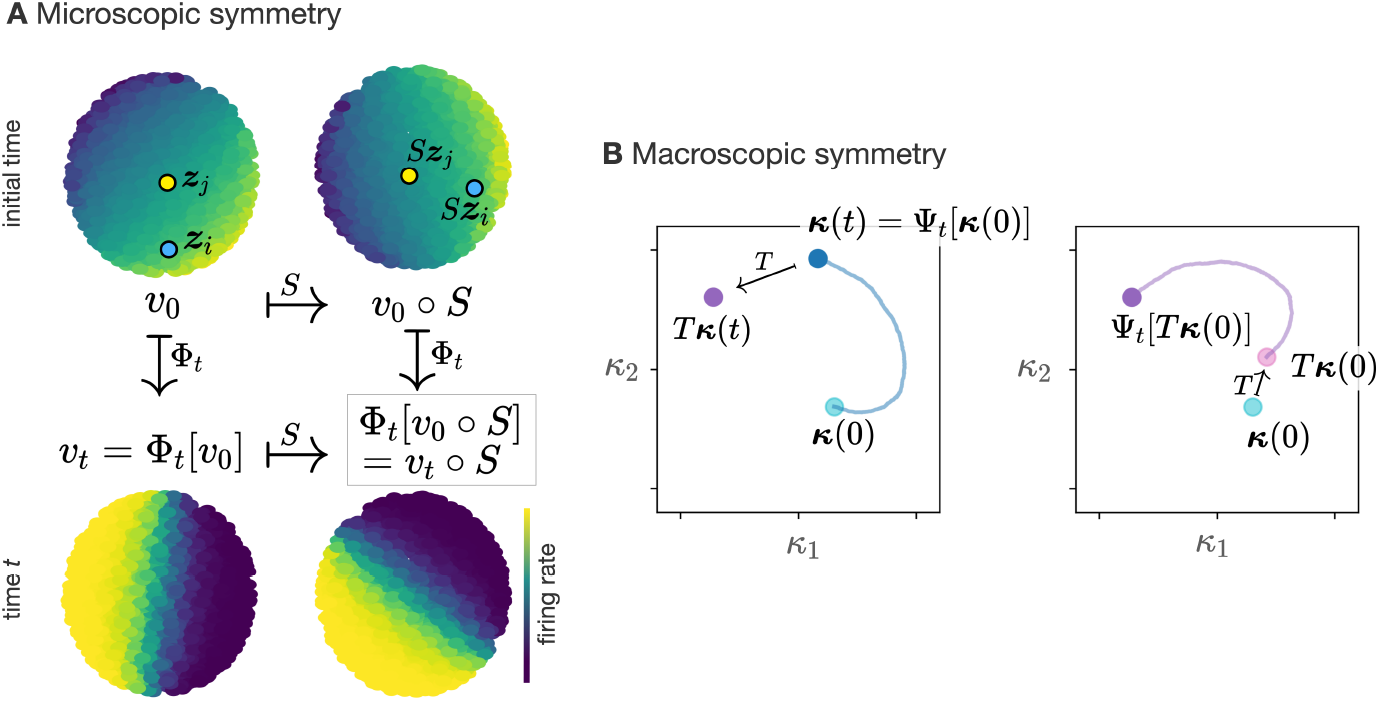
A symmetry is a mapping that commutes with the flow. **A**. Heatmaps represent the field of instantaneous firing rates, *ϕ*(*v*_*t*_(*z*)), over a disc-like, *d* = 2-dimensional similarity space, at initial time (upper panels) and final time *t* (lower panels). A *microscopic symmetry* of a field model is a transformation *S* : *ℳ* → *ℳ* of the neuronal locations that commutes with the flow Φ_*t*_ of the field dynamics up to any time *t*, i.e. such that the upper and lower path are equivalent. The upper path (right-down) first composes the initial field *v*_0_ with *S*, then applies Φ_*t*_, yielding the final state Φ_*t*_[*v*_0_ ° *S*]. The lower path (down-right) first appliesΦ_*t*_, then composes the field with S, yielding the final state *v*_*t*_ ° *S* =Φ_*t*_[*v*_0_] ° *S*. **B**. A *macroscopic symmetry* of the model is a mapping *T* : ℝ^*D*^ → ℝ^*D*^ of the latent variables that commutes with the flow Ψ_*t*_ of the collective dynamics up to any time *t*: the final state (purple dot) is the same in the left panel (first apply Ψ_*t*_, and then *T*) and in the right panel (first apply *T*, and then Ψ_*t*_). Solid lines represent the trajectory of the latent variables in a *D* = 2-dimensional neural manifold, from initial time up to time *t*, starting from the initial condition ***κ***(0) (left panel) or *T* ***κ***(0) (right panel).

### Symmetry groups

The mapping *𝒯* : *S* ↦ *T* ^(*S*)^ defines the action on ℝ ^*D*^ of any group *G* of *microscopic* symmetries of the model, which will also be a group of *macroscopic* symmetries. Yet, the order of composition of group elements is reversed: for any elements *S, S*^*′*^ of *G* acting on ℳ, the action on ℝ ^*D*^ of their composition *S ∘ S*^*′*^ is given by: 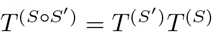. Therefore, a “right” group action on ℳ becomes a “left” group action on ℝ ^*D*^ (Corollary 1 in Appendix C). This will have an effect if the symmetry group *G* is non-commutative; like, for example, the group of rotations in ℝ ^3^.

## 4 Extracting the similarity space

### 4.1 Models of limit-cycle dynamics

As a first illustration of our method for extracting the similarity space from the neuronal activity, we studied two spiking models that generate similar limit-cycle dynamics. SNNs were implemented as in the examples of Fig. 3C-D, with connectivity given by Eqs.(8) and (9); the only difference is that we now consider a sigmoidal transfer function *ϕ*^1^. Since both models yield similar collective dynamics (Appendix B.2), the activity of the two networks generates similar low-dimensional trajectories, that both enter into a limit cycle in a *D* = 2-dimensional neural manifold (Fig. S2A,B).

To distinguish the two networks from each other, we characterised the spiking activity of each neuron with two statistical measures, chosen to be the neuron’s tuning to each of the two first PCs of the low-passed spiking activity. This yields an two-dimensional embedding of each neuron. The point-cloud of the set of all neuronal embeddings enables us to visualise the functional similarity space (up to an isomorphism) and to distinguish between the two different models (Fig. S2C): for the first SNN, the similarity space is a one-dimensional ring (Fig. S2C1); for the second SNN, it is a two-dimensional surface (Fig. S2C2). For both models, the empirical location of the neurons in the similarity space was then obtained by parameterising their relative positions in this point-cloud. A representation of the neuronal activity as a function of the empirical neuronal locations highlights the underlying spatial structure, which differs between the two models (Fig. S2D). Finally, sorting the neurons with respect to their empirical locations makes oscillatory dynamics clearly visible (Fig. S2E), similar to a recent analysis that revealed slow oscillatory dynamics in the mouse MEC [22]. We stress that sorting the neurons is equivalent to arranging them in one-dimensional space; higher-dimensional neuronal embeddings can thus be seen as a generalisation of neuron sorting (Fig. S2,D2).

### 4.2 Models of context-dependent decision-making

We reformulate the task in our framework. In a circuit model, the input weights, i.e., the contribution of the sensory and contextual signals to the external input received by each neuron *i*, are parameterised by functions of the neuron’s location ***z***_*i*_, as expressed by Eq.(2). Similarly, the output is a one-dimensional readout of the spiking activity, with weights determined by the neurons’ locations.

For both models, the contribution of the sensory and contextual input features (respectively: *I*_1_, ‘color’, *I*_2_, ‘motion’, and *I*_c_, “context”) to the external input received by each neuron *i* is parameterised by functions 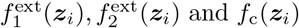 respectively. We impose the recurrent and the sensory input to be colinear by setting 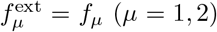. The total external input received by each neuron *i* reads:

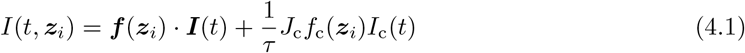

where ***I*** = (*I*_1_, *I*_2_) is the sensory input, and *J*_c_ is the strength of the context input. During each trial, *I*_1_(*t*) and *I*_2_(*t*) fluctuate around a fixed mean ±*Ī*, which is positive or negative with fixed magnitude *Ī*: i.e. *I*_*µ*_(*t*) = ±*Ī*+ *σξ*_*µ*_(*t*), where *ξ*_*µ*_(*t*) is a Gaussian white noise and *σ*^2^ is the variance of the fluctuations. The context input is unitless and takes binary values *I*_c_(*t*) = ± 1 during trials; and the factor 1/*τ* takes care of the units.

Since the maps *f*_1_, *f*_2_, *f*_*c*_ determine the contribution of the sensory features *I*_1_, *I*_2_ and context *I*_*c*_ to the external input each neuron receives, they can be understood as encoding the neuron’s selectivity (or tuning) to these features.

#### Collective dynamics

We assume symmetric couplings, i.e. ***g*** = *J****f*** ; and arbitrary voltage-to-rate function *ϕ*. For a stationary context input *I*_c_(*t*) = ± 1, the collective dynamics write (from their general expression, Eq.(A.13)):

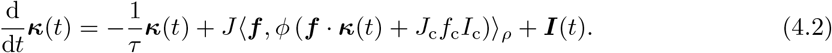

**Fig. S2.**
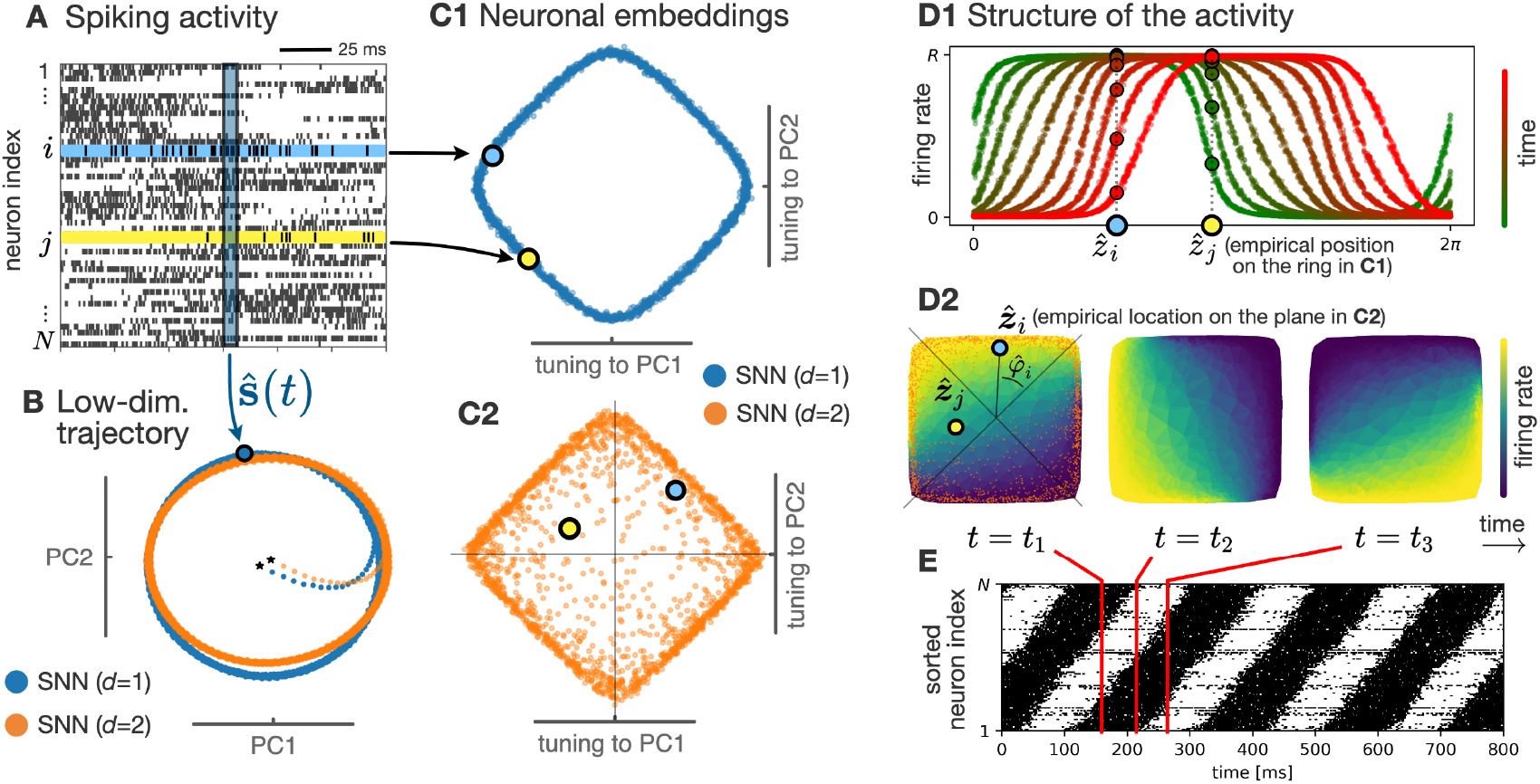
The functional similarity space can be extracted from the neuronal activity. Two SNNs of *N* = 2000 neurons follow similar limit-cycle collective dynamics, even though they have a different cicuit structure (Fig. 3C,D). **A**. Spike trains of randomly chosen example neurons in the SNN with *d* = 1, with two neurons *i* and *j* highlighted. At any time *t*, the *N*-dimensional population activity lies on the neural manifold. In contrast, the spike train of each neuron *i* depends on its location ***z***_*i*_ in the similarity space. **B**. The trajectory of the two SNNs is shown in the space of the first two principal components (PCs) of the low-passed spiking activity 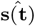 (> 80% variance explained for both networks), after a rotation to superimpose the two trajectories. Both trajectories enter a limit-cycle with the same period *T* = 195 ms. **C**. The spiking activity of each neuron was characterised by its tuning to each of the two first principal components (PCs) of the population activity. **C1**. For the first SNN, the point-cloud formed by the embeddings of all the neurons in the population forms a ring. **C2**. In constrast, for the second SNN, the set of all two-dimensional embeddings lies on a two-dimensional surface. For both networks, it is consistent with the similarity space of the underlying circuit model (Fig. 3C). **D1**. Each neuron *i* of the first SNN is assigned an empirical location 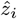 according to its position on the ring-like point-cloud of panel C1. Coloured curves represent the neurons’ firing rates as a function of their empirical locations, at several time points (where colour from green to red represents increasing time). The time series of the firing rate of two neurons *i* and *j* are highlighted. **D2**. Each neuron *i* of the second SNN is assigned a two-dimensional empirical location 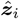 according to its position in the point-cloud of panel C2. Heatmaps represent the neurons’ momentary firing rates as a function of their empirical locations, at three time points *t*_1_, *t*_2_, *t*_3_ (from left to right). Orange dots on the left-most heatmap represent all empirical locations inferred from panel C2. The whole point-cloud is displayed with a 45° rotation compared to panel C2 for convenience. **E**. Rasterplot of the spiking activity of the second SNN, where neurons were sorted according to the angle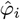 of their empirical location (panel D2, left). This makes oscillatory dynamics clearly visible, similar to the analysis performed in [22]. Red bars correspond to the three time points shown in panel D2. Details about implementation in Methods 4.3.

The connectivity functions ***f*** (*z*), the context selectivity *f*_c_(*z*) and the strength parameters *J, J*_c_ were chosen such that in the absence of external input (*I*_1_ = *I*_2_ = *I*_c_ = 0), these dynamics admit no non-trivial fixed point; but in the ‘color’ context (*I*_c_ = +1), symmetric attractive fixed points appear along the ‘color’ axis at ***κ*** = (±*k*, 0), with *k*> 0 given by the fixed points of Eq.(4.2) with *I*_c_ = +1. Similarly, the ‘motion’ context (*I*_c_ = −1) yields attractive fixed points along the ‘motion’ axis at ***κ*** = (0, ±*k*), as illustrated in Fig. 4B. More details about the implementation are given below.

This was achieved by setting *f*_c_ = |*f*_1_| − |*f*_2_|. In the ‘color’ context *I*_c_ = +1, neurons that are more selective to the first input feature (‘color’), i.e. for which |*f*_1_(***z***_*i*_) | > |*f*_2_(***z***_*i*_) |, are excited; while those that are more selective to the second input feature (‘motion’) are inhibited; and reciprocally in the ‘motion’ context *I*_c_ = −1.

At the beginning of each trial, the context input is switched on to *I*_c_ =± 1, indicating the context. The population activity is then driven towards one of the fixed points by the relevant component of the sensory input (*I*_1_, ‘color’ if *I*_c_ = +1, or *I*_2_, ‘motion’ if *I*_c_ =− 1), while ignoring the irrelevant component; and it relaxes towards the baseline state when the context input is switched off, being ready for a new trial.

#### Choice readout

The goal of the task is to match the output of the network with the sign of the relevant input feature, which is specified by the context. The output is a one-dimensional linear readout of the activity, with output weights *f*_out_(***z***_*i*_):

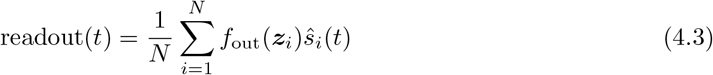

with *ŝ*_*i*_(*t*) the low-passed spike train of neuron *i*. The map of output weights is set to *f*_out_ = *f*_1_ +*f*_2_. Thus, the readout is positive for the two fixed points that correspond to a positive relevant sensory input: (+*k*, 0) in the ‘color’ context if *I*_1_ > 0, or (0, +*k*) in the ‘motion’ context if *I*_2_ > 0. Similarly, the readout is negative for a negative relevant sensory input.

#### Low-dimensional trajectory of the SNN

Low-dimensional trajectories of the SNNs in Figs.4E and 5B were obtained by projecting the low-passed *N*-dimensional spiking activity onto ‘color’ and ‘motion’-relevant axes, defined 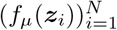 (with *μ*= 1 for the ‘color’ and *μ*= 2 for the ‘motion’ axis). Since *f*_out_ = *f*_1_ + *f*_2_, the ‘choice’ readout of Eq.(4.3) is equivalent to the projection of this 2-dimensional population response on the diagonal axis (1, 1) (Fig. 4C,E).

#### Implementation with two circuit models

The first “blue” model in the main text is a ring model: neurons have angular positions uniformly distributed over [0, 2*π*[; and the connectivity is given by *W*_*ij*_ = *J* cos(*z*_*i*_− *z*_*j*_) (as in Eq.(8), with *δ* = 0) which is written in rank-2 form using ***f*** (*z*) = (cos *z*, sin *z*).

The second “orange” model has a 2-dimensional functional similarity space: neuronal locations are uniformly distributed in the square [−1, 1]^2^ (i.e., ***z***_*i*_ = (*z*_*i*,1_, *z*_*i*,2_) where *z*_*i*,1_ and *z*_*i*,2_ are independent and uniformly distributed in [−1, 1]).The connectivity is the dot product of the locations: we set ***f*** (***z***_*i*_) = ***z***_*i*_, so that *W*_*ij*_ = *J* ***z***_*i*_ · ***z***_*j*_. For both models, the voltage-to-rate function was a shifted sigmoid: *ϕ*(*x*) = *R*(1 + tanh(*x* − 1))/2. In the absence of sensory external input, Eq. (4.2) admits fixed points given by

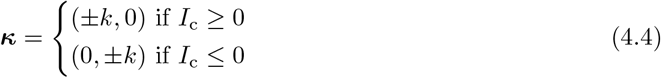

where *k* ≥ 0 is the solution to:

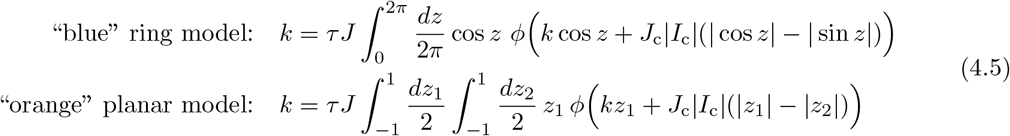

The values of the parameters *J* and *J*_c_ were chosen so that the above equations have no solutions *k* ≠ 0 for *I*_c_ = 0 (corresponding to the “baseline” dynamics in Fig. 4B), but have a non-trivial solution *k*> 0 for |*I*_c_| = 1. Solutions were found numerically. The chosen values were *J* = 0.7, *J*_c_ = 1.5 for both models (given the value of *τ R* = 10).

#### Additional task-irrelevant connectivity component

We finally consider a modification of the “blue” ring model, where each neuron is assigned an additional feature *ξ*_*i*_ = ± 1, chosen randomly for each neuron *i*, and the recurrent connectivity is given by:

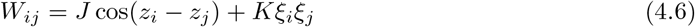

with *K*≥ 0. Each neuron *i* is thus characterised by its location on a double-ring, (*z*_*i*_, *ξ*_*i*_) ∈ [0, 2*π*[×{±1}. The recurrent connectivity has rank 3, with connectivity functions given by ***f*** (*z, ξ*) = (cos *z*, sin *z, ξ*) ∈ ℝ ^3^, and ***g***(*z, ξ*) = (*J* cos *z, J* sin *z, Kξ*). The collective dynamics are three-dimensional, and read in the absence of sensory external input (similar to Eq.(4.2)):

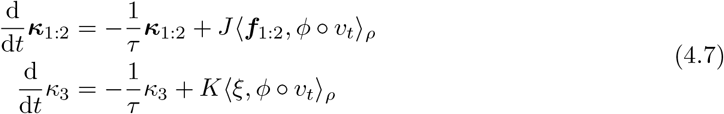

where v_*t*_(*z, ξ*) = ***f***_1:2_(*z*) · ***κ***_1:2_(*t*) + *ξκ*_3_(*t*) + *J*_c_*I*_c_*f*_c_(*z*), and ***κ***_1:2_ = (*κ*_1_, *κ*_2_), ***f***_1:2_ = (*f*_1_, *f*_2_).

Notice that if *κ*_3_ = 0, then *v*_*t*_ does not depend on *ξ*. Thus ⟨*ξ, ϕ* ° *v*_*t*_⟩_*ρ*_ = 0; and the first line of Eq.(4.7) becomes identical to Eq.(4.2). Therefore, the fixed points of Eq.(4.2) for ***κ***_1:2_ (given by Eq.(4.4)) are also fixed points of Eq.(4.7) given *κ*_3_ = 0. Yet, those fixed points should be stable (in the direction of *κ*_3_), in order for the model to behave similarly to the original “blue” model. The condition of stability in the direction of *κ*_3_ reads (from Eq.(4.7)):

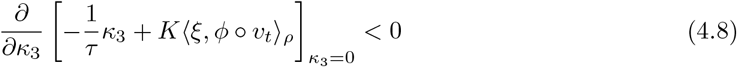

which yields, after explicit computation:

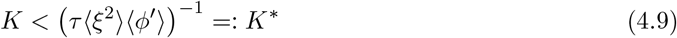

where ⟨*ξ*^2^⟩ (= 1 in our case) is the average value of *ξ*^2^; and 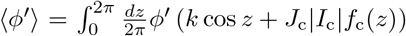 is the average value of the “gain” function *ϕ*^*′*^.

In Fig. 5, *K* was set near below the critical value *K*^∗^ at which the fixed points with *κ*_3_ = 0 lose their stability. In addition, the network received a weak external input in the direction of *κ*_3_, of the form

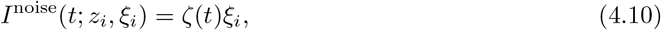

where *ζ*(*t*) is an Ornstein-Uhlenbeck process with mean 0 and small variance. In the original “blue” model with *K* = 0, the effect on the dynamics was negligible. In the model with *K* close to the critical value *K*^∗^, the additional circuit structure dramatically enhances this weak input, leading to large fluctuations of *κ*_3_, and thus to a strong dependence of the membrane potentials (and thus of the firing rates) on the task-irrelevant features *ξ*_*i*_.

The value of the critical strength (Eq.(4.9)) was computed to be *K*^∗^ ≈ 0.67 for *I*_c_ = ±1, and *K*^∗^ ≈ 0.48 for *I*_c_ = 0 (given the parameters *τ R* = 10, and *J* = 0.7, *J*_c_ = 1.5; which yields from Eq.(4.5) *k* ≈ 1.29 for *I*_c_ = ±1, and *k* = 0 for *I*_c_ = 0). *k* was then set to 0.4.

### 4.3 Numerical simulations

In all simulations (Figs. 3, 4, 5, S2), the neuronal membrane time constant was set to *τ* = 10 ms, which defines all characteristic time scales. The maximum firing rate was *R* = 10/*τ*. In Figs. 4, 5, S2, spikes were low-passed with a time constant *α* = *τ*, and the spike train of each neuron was then z-scored by substracting its mean and dividing by its standard deviation. PCA was performed on the z-scored low-passed activity.

For the decision-making task, analyses were performed on 8 trials that covered all possible combinations of the 3 binary task variables (mean ± *Ī* of the ‘color’ and ‘motion’ inputs, and value ± 1 of the ‘context’ input) (only 4 trials are shown on Figs. 4 and 5). The magnitude of the sensory inputs (see below Eq.(4.1)) was set to *Ī* = 0.1*τ* ^−1^, and their standard deviation to *σ* = *Ī*/5.

In Figs. 4 and 5, all three models described above (“blue”, “orange” and “pink”) received the same low-dimensional input, including the sensory and context features, and also the noisy input with binary random projections *ξ*_*i*_ given by Eq.(4.10). The effect on the dynamics was negligible for models whose connectivity was uncorrelated with *ξ*_*i*_ (“blue” and “orange” SNNs). Parameters of the O.-U. process *ζ*(*t*) were *θ* = 0.1*τ* ^−1^ and *σ* = *θ*/5.

In Fig. S2, the tuning of each neuron *i* to the two first PCs was computed as the normalized difference between spike counts of the neuron during periods where each PC was positive or negative:

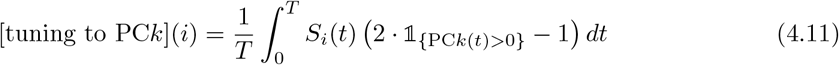

where PC*k*(*t*) (*k* = 1, 2) represents the trajectory of the *k*-th principal component of the activity, and *S*_*i*_(*t*) the spike train of neuron *i*. This choice of statistics allowed one to keep information about the variance of each neuron’s activity, which was varying across neurons for the SNN with *d* = 2. In contrast, this information was collapsed in the PC loadings associated with the z-scored population activity (due to the z-scoring).

Appendix

Note: in the Appendix, we drop the boldface notation ***z*** to denote neuronal locations. Instead, the notation *z* still represents a *d*-dimensional variable in the functional similarity space ℳ.

## A Collective dynamics

### A.1 Derivation of the collective dynamics

We start from Eq.(1.6), rewritten here for convenience:

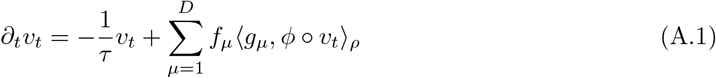

The recurrent input belongs to the linear subspace ℱ = span{*f*_*µ*_} ⊂ ℋ. We can decompose the field *v*_*t*_ as its projection onto ℱ, plus a component 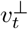 belonging to its orthogonal complement ℱ^⊥^:

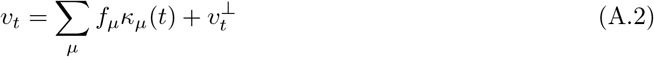

which implies:

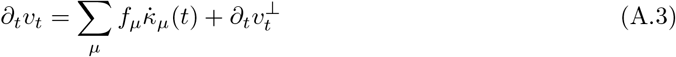

where the dot (·) denotes the time derivative. Together, equations (A.1)-(A.2) yield:

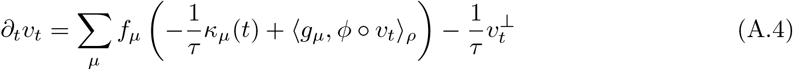

As both 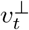 and 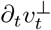 are orthogonal to all of the *f*_*µ*_, we get by identification of eqs.(A.3) and (A.4):

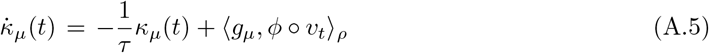

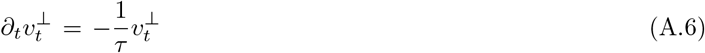

Thus, 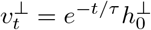, so that Eq.(A.2) yields Eq.(1.7). It allows replacing v_*t*_ in Eq.(A.5) to obtain, in vector form:

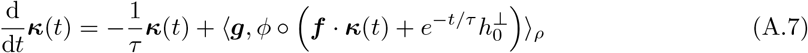

which yields Eq.(1.9) if we assume *h*_0_∈ ℱ.

Eventually, the expression of the latent variables ***κ*** can be obtained by projecting Eq.(A.2) on ℱ: ⟨*f*_*ν*_, *v*_*t*_⟩_*ρ*_ = Σ_*µ*_⟨*f*_*ν*_, *f*_*µ*_⟩_*ρ*_ *κ*_*µ*_; or in vector form: ⟨***f***, *v*_*t*_⟩_*ρ*_ = C***κ***, with the correlation matrix C_*νµ*_ := ⟨*f*_*ν*_, *f*_*µ*_⟩ _*ρ*_. We can assume without loss of generality that the set of functions {*f*_*µ*_}are linearly independent in ℋ. If that is not the case, a new basis of *ℱ* with fewer elements can always be chosen (see also the next section A.2). Thus, the matrix *C* has full rank *D*, and is thus invertible, yielding Eq.(1.8):

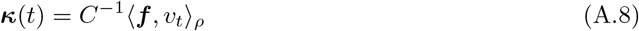

### A.2 Including external inputs

Assume that the network receives an external input composed of *K* features *I*_*ν*_(*t*) (*ν* = 1, …, *K*), to each of which the tuning of any neuron *i* is determined by a corresponding function 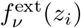 of its location *z*_*i*_. The external input received by any neuron can be written: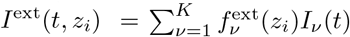. The total (recurrent + external) input thus now belongs to the linear subspace 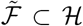 spanned by 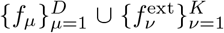, of dimension 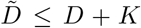. Let 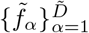 be a basis of 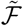. Since any *f*_*µ*_ ∈ *ℱ* is a linear combination of the 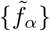,then for each α, there exists a linear combination 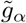 of the feature maps {*g*_*µ*_} such that

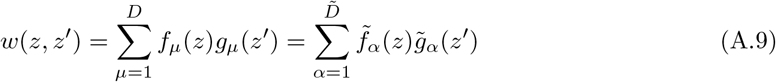

and similarly, there exists a linear combination *Ĩ*_*α*_ of the input features {*I*_*ν*_} such that

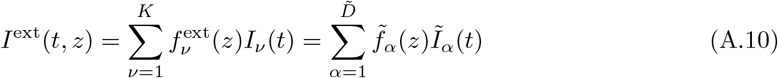

Without loss of generality, we can thus write the total synaptic input received at location *z* as:

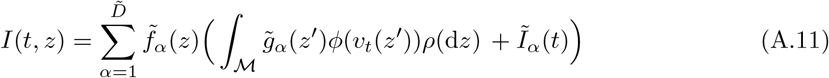

From now on, we omit the tildes 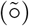 for readability. The field equation in the case of a low-dimensional external input can be written:

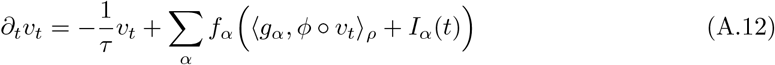

Repeating the derivation of Sec.A.1 gives the general expression of the collective dynamics:

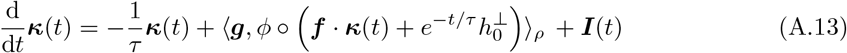

where ***I***(*t*) = (*I*_1_(*t*), …, *I*_*D*_(*t*)) is the vector of external input features.

### A.3 Generalisation to arbitrary square-integrable kernels

We turn now to the case of field models where the connectivity is determined by a kernel *w*(*z, z*^*′*^) over ℳ^2^ which is only assumed to be in *L*^2^(*ρ* × *ρ*), i.e.

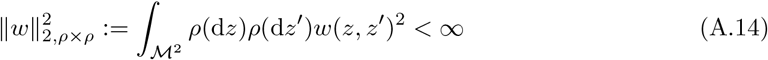

yielding the general field equation:

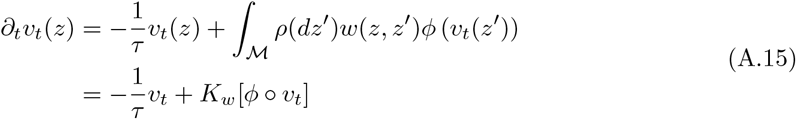

where *K*_*w*_ : ℋ → ℋ is the Hilbert-Schmidt integral operator associated with the kernel *w*: [92]

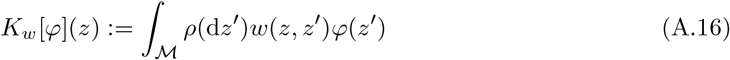

Since *K*_*w*_ is a Hilbert-Schmidt operator, it can always be decomposed as:

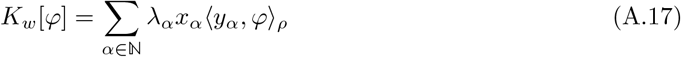

where the sets of left and right eigenvectors {*x*_*α*_}, {*y*_*α*_} are orthonormal families in ℋ. The l^2^-norm of the series of singular values λ_*α*_ ≥ 0 is finite, and is equal to the Hilbert-Schmidt norm of i.e. 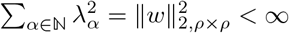 [92, 93]. The dynamics of general fields, Eq.(A.15), have thus a very similar structure to that of low-rank fields, Eq.(1.5) – except (precisely) that the rank of *K*_*w*_ is not necessarily finite. In particular, one can still write the associated collective dynamics for the projections *κ*_*α*_(*t*) = ⟨x_*α*_, u_*t*_⟩_*ρ*_, following the line of sec.A.1:

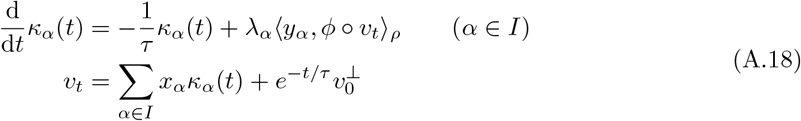

where *I* = {*α* | *λ*_*α*_ > 0} ⊆ ℕ, and 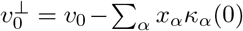 From this point of view, a low-rank field is only characterised by a finite set of non-zero singular values λ_*α*_ > 0. The dynamics of any arbitrary field can be approximated using a truncation of the sum in Eq.(A.17) [59], which corresponds to a model of low-rank network according to our framework (see Eq.(1.3)).

## B Examples of collective dynamics

### B.1 Fixed points in a generalised Hopfield model

We consider here the connectivity mappings given by Eq.(2.2), which read:

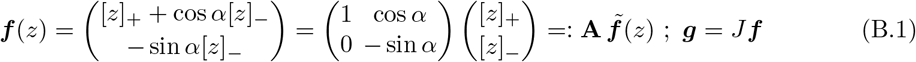

with 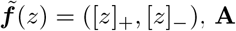 the 2 × 2 matrix above, and [*z*]_*±*_ = *z* · 𝟙_{sgn(*z*)=*±*1}_. Let the neurons be normally distributed over the real line, i.e. *z*_*i*_ ∼ *𝒩*(0, 1). The collective dynamics of Eq.(1.9) write:

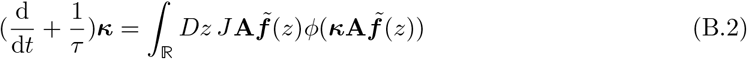

with *Dz* the standard Gaussian measure; and we assume for now a general transfer function *ϕ*. We make the following change of variable: ***λ*** = **A**^*T*^ ***κ***. Noticing that 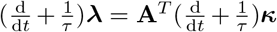, we have:

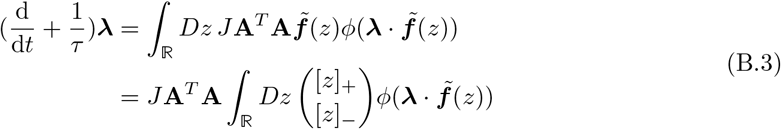

Notice that:

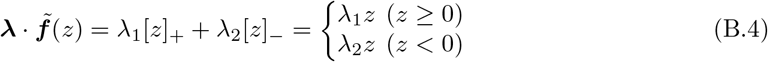

so that Eq.(B.3) can be written, by splitting the integral over ℝ :

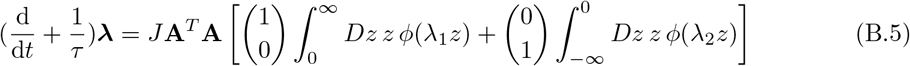

We now derive further the dynamics for the step transfer function *ϕ*(*x*) = *R*𝟙_{*x>*0}_. Notice that (i) for *z*> 0, *λ*_1_*z*> 0 ⇔ λ_1_ > 0; and (ii) for *z* < 0, λ_2_*z*> 0 ⇔ λ_2_ < 0. The above gives:

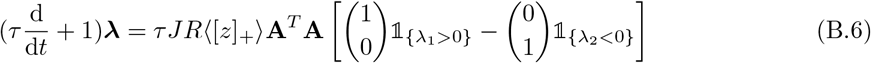

Where 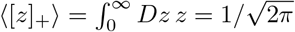. Let us define *k* = *τ JR*⟨[*z*]_+_⟩, and:

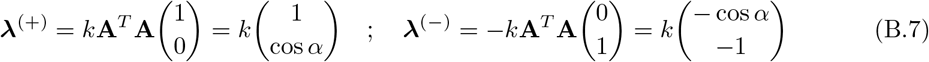

We can write:

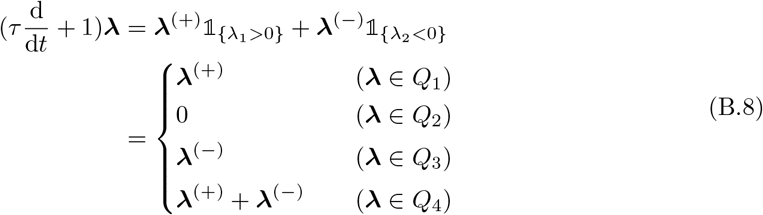

with *Q*_1≤*i*≤4_ the four quadrants of ℝ ^2^, numbered in trigonometric order. For α ∈]0, *π*/2[, ***λ***^(+)^∈*Q*_1_, ***λ***^(−)^∈ *Q*_3_ and (***λ***^(+)^ + ***λ***^(−)^)∈*Q*_4_. They are thus stable fixed points of the dynamics, while ***λ*** = 0 is a saddle point. The dynamics of the original variables ***κ***, shown in fig.3(B), are obtained through the change of coordinates ***κ*** = (**A**^*T*^)^−1^***λ***. In particular, the three corresponding fixed points are:

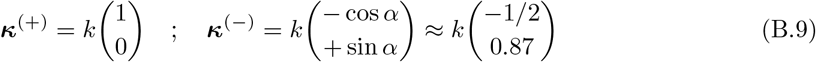

and (***κ***^(+)^ + ***κ***^(−)^). The numerical values correspond to Fig.3(B), where *α* = *π*/3 (i.e. *a* = cos *α* = 1/2).

### B.2 Different models of limit-cycle collective dynamics

#### Ring model

For the ring model, we can write from Eq.(8):

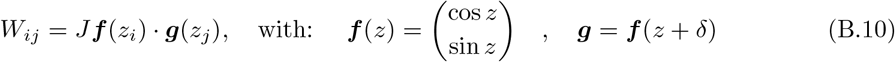

and the collective dynamics resulting from Eq.(1.9) write:

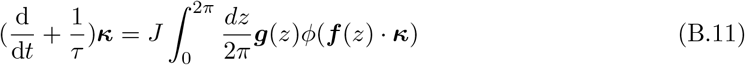

Let *ϱ, θ* be the polar coordinates of ***κ***. In the polar coordinate system (*ê*_*ϱ*_, *ê*_*θ*_) associated with ***κ***, we have:

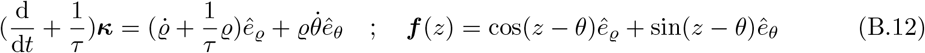

hence:

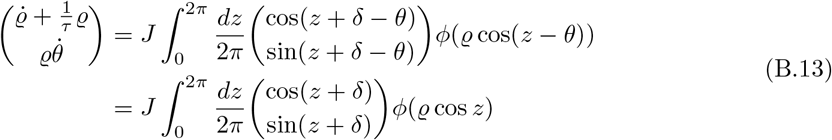

For the step transfer function *ϕ*(*x*) = *R*𝟙_{*x>*0}_, we have *ϕ*(*ϱ* cos *z*) = *R*𝟙_{−*π/*2*<z<π/*2}_, hence:

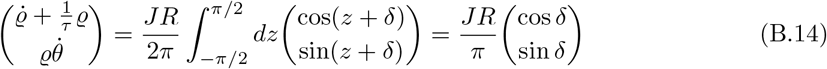

which is identical to Eq.(2.5).

### Gaussian distribution of neuronal locations

Let the neurons be normally distributed in ℝ ^2^, i.e. *z*_*i*_∼ *𝒩* (0, 𝟙_2_). As discussed in Methods 2, this distribution can be parameterised in polar coordinates: each neuron is independently assigned a radius *ξ*_*i*_ ≥ 0 and an angle*φ*_*i*_ ∼ 𝒰([0, 2*π*[). The connectivity of Eq.(9) thus writes:

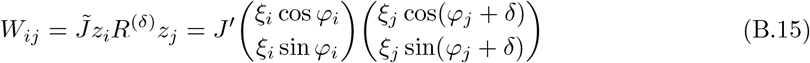

which can be written, as a function of a neuron’s radius *ξ* and angle*φ*:

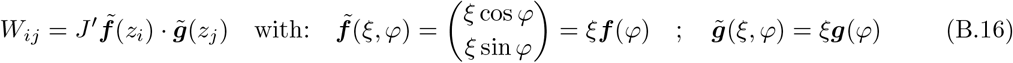

where ***f***, ***g*** are defined by Eq.(B.10). The collective dynamics of Eq.(1.9) thus write:

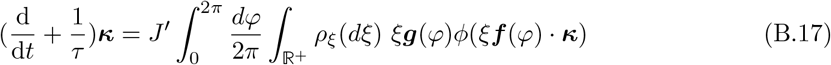

where *ρ*_*ξ*_(*d*ξ*) denotes the distribution of the radii (it is the χ_2_ distribution if neuronal locations are normally distributed in the plane). In the case of a step transfer function *ϕ*, we have *ϕ*(*ξ*x*) = *ϕ*(*x*) ∀*ξ*> 0, hence:

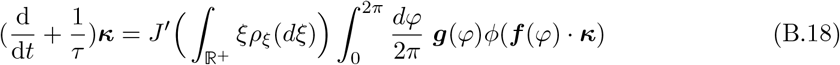

which is identical to Eq.(B.11) if *J*^*′*^⟨*ξ*⟩ = *J*, where ⟨*ξ*⟩ := *ξρ*_*ξ*_(*dξ*) (= *π*/2 for the χ_2_-distribution). Hence, the two models’ collective dynamics are identical up to a rescaling of the connectivity.

Notice that the derivation is not restricted to a normal distribution over ℝ ^2^, but applies to any rotationally invariant distribution; in this case, the only difference will be the distribution of the radii *ξ*_*i*_, which will affect the rescaling factor ⟨*ξ*⟩.

#### Arbitrary transfer function and radial distribution

Here, we extend the above derivation of the collective dynamics for the connectivity given by Eq.(B.15) to the case of an arbitrary transfer function *ϕ* and rotationally invariant distribution of the neuronal locations. In this case, the collective dynamics (Eq.(1.9)) corresponding to the connectivity functions of Eq.(B.16) are:

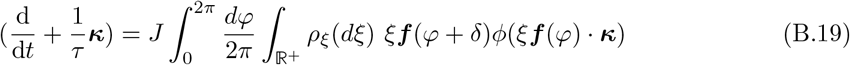

where ***f*** = (cos, sin) (Eq.(B.10)), and *ρ*_*ξ*_(*dξ*) is the distribution of the radii of neuronal locations. For the radial coordinate (*ϱ, θ*) of ***κ***, it writes:

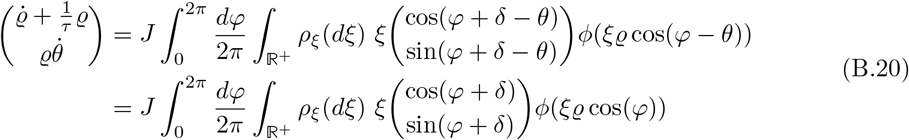

Using the sum expansion of the cos and sin functions, we get for the first component of Eq.(B.20):

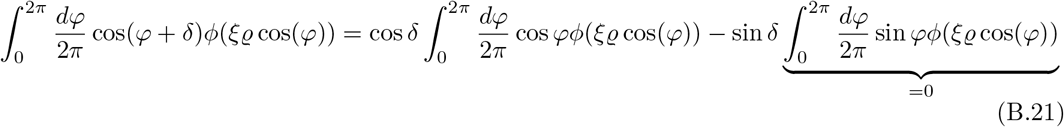

where the last term is zero because the integrand is odd; and similarly, for the second component:

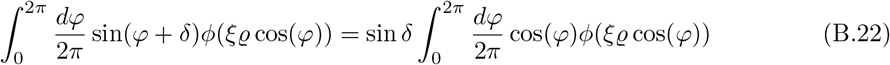

Thus, we obtain from Eq.(B.20):

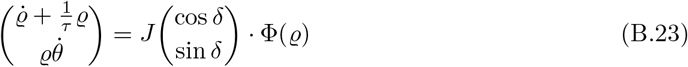

where Φ (*ϱ*) is defined by (after the change of variable *x* = cos*φ*):

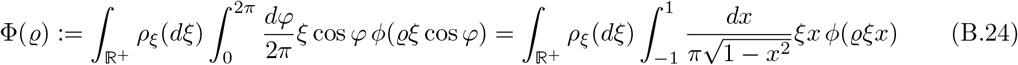

Provided that the fixed point equation for *ϱ*,

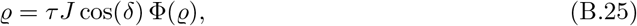

admits a solution *ϱ*^∗^, the model generates rotational dynamics that converge towards a limit cycle of radius *ϱ*^∗^.

Noticeably, the angular speed *ω* of the limit cycle does not depend on the transfer function *ϕ*, nor on the distribution *ρ*_*ξ*_ of the radii, but only on the parameters *τ* and *δ*. From Eq.(B.23) and using the definition of *ϱ*^∗^ (Eq.(B.25)), we get

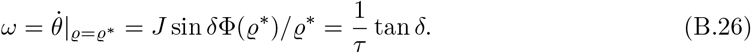

## C Symmetries

We show here the results of Sec.2, relating the symmetries of a field model to those of its collective dynamics. We take an operator theory perspective. The dynamics of the field, Eq.(1.6), can be written:

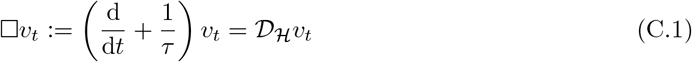

where 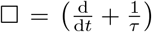 is linear, and *D*_*H*_ : ℋ → ℱ (where ℱ = span{*f*_*µ*_}) is a nonlinear operator defined by:

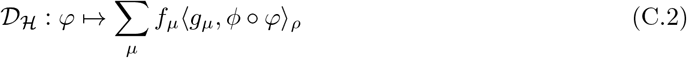

On the other hand, the collective dynamics of Eq.(1.9) are described by a nonlinear operator *D* _*D*_ acting on ℝ ^*D*^:

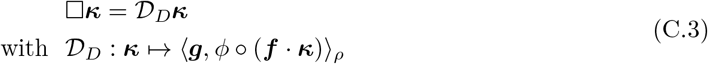

### Defining symmetries

In the following, let 𝒮 be the set of bijections *S* : ℳ → ℳ such that *φ* ° *S* ∈ ℋ, ∀*φ* ∈ ℋ. For any mapping *S* ∈ *𝒮*, the composition operator *L*^(*S*)^ :*φ* ⟼ *φ* ° *S* defines a linear operator on ℋ.

We call a *(microscopic) symmetry* of a field model a bijection *S* ∈ *𝒮* such that the operator *𝒟* _*ℋ*_ describing the field dynamics commutes with *L*^(*S*)^; i.e. *𝒟* _*ℋ*_° *L*^(*S*)^ = *L*^(*S*)^ ° *𝒟*_*ℋ*_. It implies the equivariance of the field dynamics for *L*^(*S*)^: if *v*_*t*_ solves the dynamics of Eq.(C.1), then *L*^(*S*)^*v*_*t*_ is also a solution to the dynamics, with initial condition *L*^(*S*)^*v*_0_. Analogously, a *macroscopic* symmetry is a symmetry of the collective dynamics; i.e., a mapping *T* : ℝ ^*D*^ → ℝ ^*D*^ that commutes with *D*_*D*_.

### Results

We show that any microscopic symmetry *S*∈ 𝒮 of the field model generates a macroscopic symmetry *T* ^(*S*)^ of its collective dynamics, for which we give an explicit expression – under an additional condition that is satisfied by invariance symmetries.

#### Theorem 1.

*Let S ∈ 𝒮*.

1. *If (i) S is a symmetry of the field model, and (ii)* ℱ = span {*f*_*µ*_} *is an invariant subspace of L*^(*S*)^, *then a symmetry of the collective dynamics is given by the linear operator T* ^(*S*)^ : ℝ ^*D*^→ ℝ ^*D*^, *defined by the matrix*

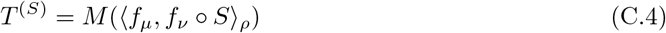

*where* ℳ ^−1^ = (⟨ *f*_*µ*_, *f*_*ν ρ*_⟩), *and* (*A*_*µν*_) *denotes the D* × *D matrix with entries A*_*µν*_.
2. *If both the distribution ρ and the connectivity kernel w*(*z, z*^*′*^) = ***f*** (*z*) · ***g***(*z*^*′*^) *are invariant for S, then the conditions (i) and (ii) of point 1 are satisfied*.

The proof can be found at the end of this section.

### Left and right symmetry groups

Let (*g, z*) ⟼ *S*_*g*_(*z*) denote the action of a group *G* on ℳ. It is called a *left* group action if *S*_*gh*_ = *S*_*g*_ ° *S*_*h*_, i.e. *S*_*gh*_(*z*) = *S*_*g*_(*S*_*h*_(*z*)), ∀*g, h* ∈ *G* (and a *right* group action if *S*_*gh*_ = *S*_*h*_ ° *S*_*g*_). *G* is a *symmetry group* of the field model if for any *g*∈ *G, S*_*g*_ is a symmetry. It is a *left* symmetry group if it has a left group action on ℳ (and respectively, a *right* symmetry group if it has a right group action).

Similarly, a left (or right) symmetry group of the collective dynamics is a group with a left (resp., right) action on ℝ ^*D*^, whose action of any element is a symmetry of the collective dynamics.

#### Corollary 1.

*Let G be a* left *symmetry group of the field model, such that the action S*_*g*_ *of each element g* ∈ *G on* ℳ *satisfies the requirements of Theorem 1*.*1. Then, G is a* right *symmetry group of the collective dynamics, with a linear group action on* ℝ ^*D*^ *given by* 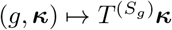.

**Proofs**.

Let *S* ∈ 𝒮; and *L*^(*S*)^ : ℋ → ℋ :*φ* ⟼*φ* ° S be the linear composition operator on ℋ.

We start by defining the surjection *K* : ℋ → ℝ ^*D*^ that maps a field to the components of its projection on ℱ, and the reciprocal injection *V* : ℝ ^*D*^ → ℱ ⊂ ℋ:

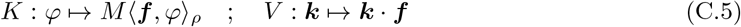

with ℳ ^−1^ = (⟨*f*_*µ*_, *f*_*ν*_⟩_*ρ*_) the inverse correlation matrix. They satisfy the following properties:

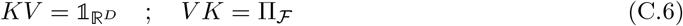

where Π_*ℱ*_ :*φ* ↦ Σ_*µ,ν*_ ℳ_*µν*_*f*_*µ*_⟨*f*_*ν*_,*φ*⟩_*ρ*_ is the orthogonal projection on ℱ (by definition of the matrix ℳ). Thus, *K* : *v* ↦ ***κ*** and *V* : ***κ*** ↦ *v* establish a bijection between ℱ and ℝ ^*D*^.

We start with a few useful lemmas.

#### Lemma 1.

*𝒟*_*D*_ = *K𝒟*_*ℋ*_*V*.

*Proof*. By setting ***κ***(*t*) = *Kv*_*t*_ and *v*_*t*_ = *V****κ***(*t*), and noticing that □ = (∂_*t*_ − 1/*τ*) commutes with *K*, one obtains from Eq.(C.1) the following expression for the dynamics of ***κ***:

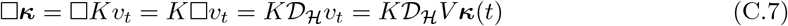

which can be identified with Eq.(C.3)).

#### Lemma 2.

*If* ℱ *is an invariant subspace of an operator L on ℋ, then LV* = Π_*ℱ*_ *LV*.

*Proof*. For any*φ* ∈ ℋ, *Vφ* ∈ ℱ. ℱ is an invariant subspace of *L* iff *Lv* ∈ ℱ ∀*v* ∈ ℱ. Then ∀*φ* ∈ ℋ, *LVφ* ∈ ℱ, implying Π_*ℱ*_ *LVφ* = *LVφ*.

#### Lemma 3.

*If ρ is invariant for the bijection S, then* 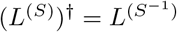

*Proof*. 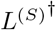 is defined by: 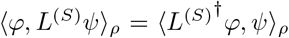 for all *φ, ψ* ∈ ℋ. If *ρ* is invariant for *S*, i.e. *ρ* ° S^−1^ = *ρ*, then for all *φ, ψ* ∈ ℋ,

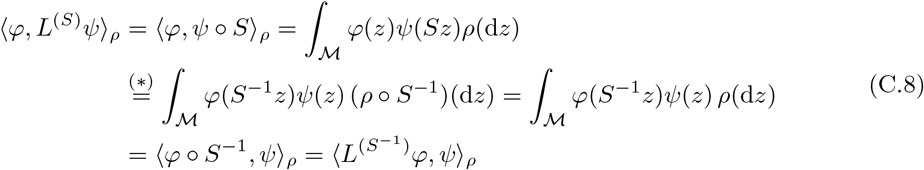

where we used a change of variable in (∗).

***Proof of Theorem 1.***

***Point 1***.

Let *L* be a linear operator on ℋ that satisfies: (i) *𝒟*_*ℋ*_*L* = *L𝒟*_*ℋ*_, and (ii) ℱ is an invariant subspace of *L*. We have from (i):

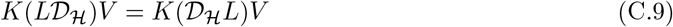

The left-hand side can be written:

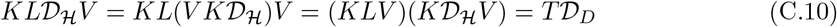

with *T* := *KLV*. The first equality is obtained by noticing that the image of *𝒟*_*ℋ*_ is in ℱ, so that Π_*ℱ*_ *𝒟*_*ℋ*_ = *𝒟*_*ℋ*_; combined with *V K* = Π_*ℱ*_, Eq.(C.6); and we used Lemma 1 in the last equality.

In turn, the right-hand side writes:

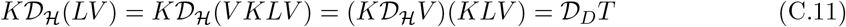

where we used Lemma 2, with *V K* = Π_*ℱ*_.

The operator *T* = *KLV* thus commutes with the operator *𝒟*_*D*_: it is a symmetry of the collective dynamics. Given a transformation *S* such that *L* = *L*^(*S*)^, we can write, for all ***κ*** ∈ ℝ ^*D*^:

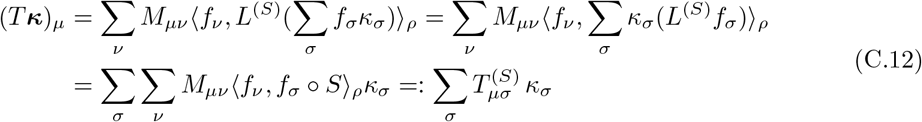

which 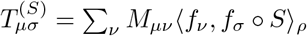. We used the linearity of L^(*S*)^ in the second equality.

***Point 2***.

Let both *ρ* and the connectivity *w* be invariant for *S* (i.e. *w*(*z, z*^*′*^) = *w*(*Sz, Sz*^*′*^) ∀*z, z*^*′*^ ∈ ℳ). We first show that it implies: *𝒟*_*ℋ*_*L*^(*S*)^ = *L*^(*S*)^*𝒟*_*ℋ*_. For all *v* ∈ ℋ and *z* ∈ ℳ, we have:

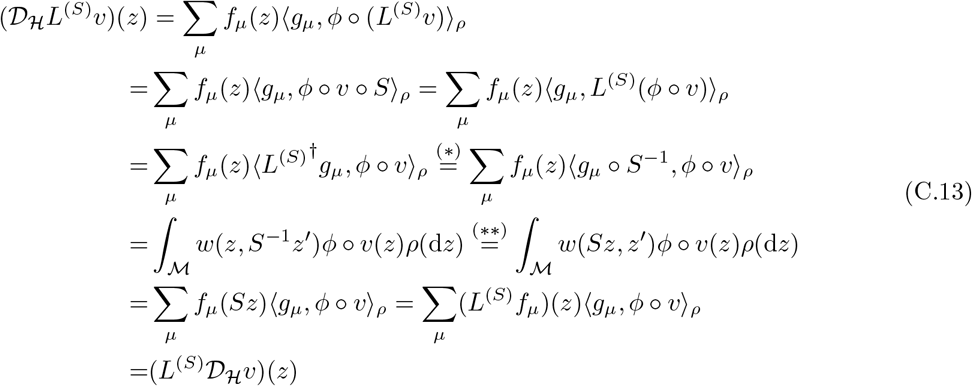

where we used Lemma 3 in (∗), and the invariance of *w* in (∗∗).

We now show that it implies that ℱ is invariant under *L*^(*S*)^. For this, it is sufficient to show that the images of the basis functions *L*^(*S*)^*f*_*µ*_ ∈ *ℱ*, ∀*μ*= 1, …, *D*. We have:

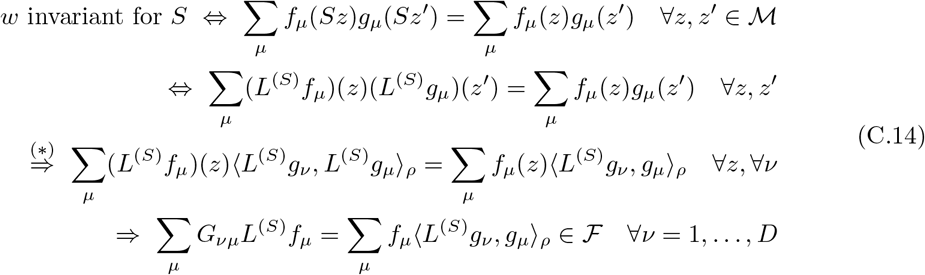

where (∗) comes by projection of the functions of *z*^*′*^ on *L*^(*S*)^*g*_*ν*_; and *G*_*νµ*_ := ⟨*L*^(*S*)^*g*_*ν*_, *L*^(*S*)^*g*_*µ*_⟩_*ρ*_ =⟨*g*_*ν*_, *g*_*µ*_⟩_*ρ*_ from the invariance of *ρ* and Lemma 3.

Without loss of generality, one can assume that the functions {*g*_*µ*_} are linearly independent. If not verified, one can always choose a new basis of ℱ such that this is the case. Thus, the matrix *G* has full rank, and is therefore invertible. The last line of Eq.(C.14) yields:

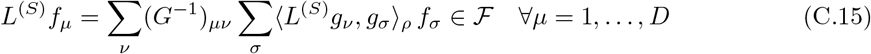

i.e. L^(*S*)^*f*_*µ*_ ∈ ℱ, ∀*μ*= 1, …, *D*, which concludes the proof.

***Proof of Corollary 1***.

In the following, we use the shorthand notation 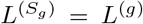 and 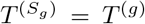 To prove the corollary, it is sufficient to show that *T* ^*(gh)*^ = *T*^*(h)*^*T*^*(g)*^,, ∀*g, h* ∈ *G*; and that 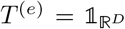 for the identity *e* ∈ *G*.

Given *G* has a left group action, we have *S*_*gh*_ = *S*_*g*_° *S*_*h*_; and thus, 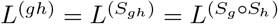. We have by associativity of the composition: 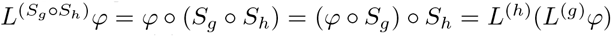, for all ℋ ∈*φ* ; hence: *L*^(*gh*)^ = *L*^(*h*)^*L*^(*g*)^. Thus, (*g,φ*) ↦ *L*^(*g*)^*φ* defines a *right* group action of *G* on ℋ.

By recalling *T* ^(*S*)^ = *KL*^(*S*)^*V*, we get:

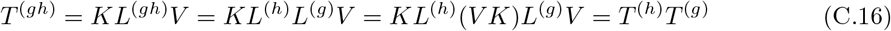

where we used Lemma 2 in the third equality, with *V K* = Π_*ℱ*_ (Eq.(C.6)).

Notice eventually that the action on ℝ ^*D*^ of the identity element *e* ∈ *G* is the identity: we have 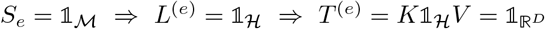 according to Eq.(C.6).

## D. Adding heterogeneity in a field model

### Rescaling of outgoing weights

Consider a field model, obeying the collective dynamics of Eq.(1.9), in which the outgoing weights of each neuron *i* are rescaled by an heterogeneity parameter *ξ*_*i*_ > 0, drawn i.i.d. across neurons according to an arbitrary distribution *ρ*_*ξ*_. This results in a “new” field model, where each neuron *i* is also characterised by its heterogeneity parameter *ξ*_*i*_, corresponding to an additional dimension of the similarity space. We denote each neuron’s location in the “new” model

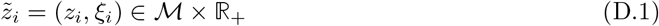

and the “new” connectivity reads:

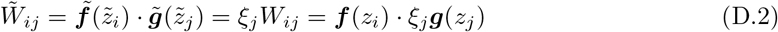

which allows us to identify 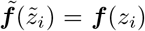, and 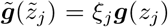.The collective dynamics, Eq.(1.9), of the latent variables 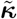 associated with this “new” model with heterogeneous connectivity write:

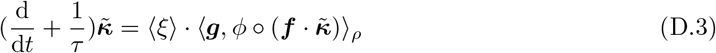

with ⟨*ξ*⟩ :=∫ *ξρ*_*ξ*_(*dξ*) the mean value of the heterogeneity parameter *ξ*. This is identical to the dynamics of the original model (Eq.(1.9)), up to a rescaling of the latent variables by a factor ⟨*ξ*⟩.

### Heterogeneous maximum firing rates

In a similar fashion, starting from an original field model, each neuron’s firing rate can be rescaled by a parameter *a*_*i*_ > 0 drawn according to a distribution *ρ*_*a*_. The firing rate of any neuron *i*, given its membrane potential *h*_*i*_, is now given by *r*_*i*_ = *a*_*i*_*ϕ*(*h*_*i*_). The transfer function 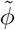 of the “new” model thus depends on each neuron’s “new” location 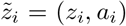: we write 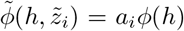.Similar to the example above (“Rescaling of outgoing weights”), the collective dynamics of the “new” and of the original field models are identical, up to a rescaling by the mean value of the heterogeneity parameter, ⟨*a*⟩ = ∫*aρ*_*a*_(*da*).

## E Field models as universal approximators

### E.1 Gaussian mixture low-rank networks

Gaussian mixture low-rank networks are a class of low-rank RNNs recently introduced to study how the collective dynamics are shaped by the statistics of the connectivity vectors [49, 60]. In this model, the rank-*D* interaction between neurons *i, j* is given by ***m***_*i*_· ***n***_*j*_, where the connectivity vectors ***m***_*i*_, ***n***_*i*_ ∈ ℝ ^*D*^ associated with each neuron *i* are drawn according to the Gaussian mixture of *P* populations:

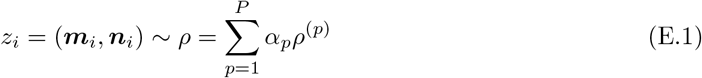

where for each *p* = 1 …, *P, ρ*^(*p*)^ is a multivariate Gaussian over ℝ ^2*D*^; and 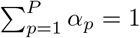. Here, each neuron’s location *z*_*i*_ ∈ ℝ ^2*D*^ corresponds to the concatenation of the connectivity vectors, and one can directly identify ***f*** (***m, n***) = ***m, g***(***m, n***) = ***n***.

These RNNs have been shown to be universal approximators of arbitrary *D*-dimensional collective dynamics, if the number *P* of populations can be chosen arbitrarily [60].

### E.2 Field models over one-dimensional space

For the sake of simplicity, we focus on the case of bounded one-dimensional space ℳ = [0, 1], with uniform distribution *ρ*(d*z*) = d*z*. We furthermore restrict to the subspace of connectivity functions *f*_*µ*_ ∈ ℋ that vanish at the boundary, that we denote ℋ_0_.

First of all, biases can be introduced by adding a constant latent variable *κ*_0_(*t*) = 1, associated with a “bias” function *f*_0_(*z*). Let the set of approximating functions, called AF_0_, be the set of functions 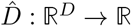 of the form

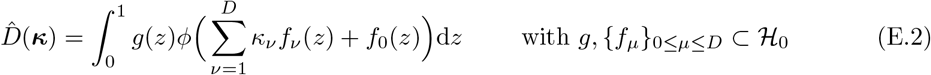

AF_0_ is a subset of the linear readouts that can be generated by field models over the one-dimensional similarity space ℳ = [0, 1].

Consider an arbitrary dynamical system with asymptotic exponential decay:

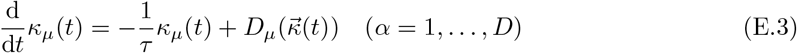

where *D*_*µ*_ ∈𝒞(ℝ ^*D*^, ℝ) vanish for 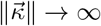. The approximation problem consists in finding functions *g* and {*f*_*µ*_} so that each given *D*_*µ*_ is approximated by some 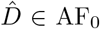. As for feed-forward neural networks, this can be decomposed into two parts [94]. First, show that approximating functions can be linearly combined: for any 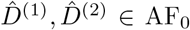 and *c*_1_, *c*_2_ ∈ ℝ, one can find 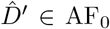 such that 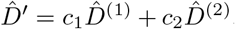. Second, show that elementary functions (like half-plane indicators) can be well approximated. Arbitrary functions can then be approximated using linear combinations of elementary functions.

#### Linear combination

For any linear combination in AF_0_, we can write:

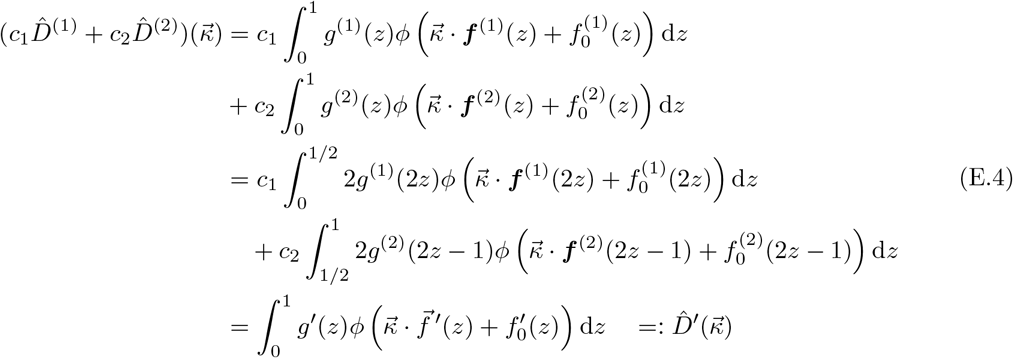

where the parameters of 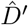 are

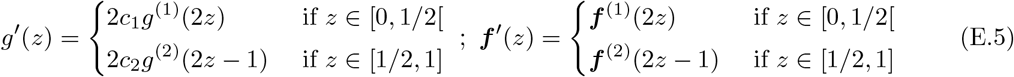

for α = 0, …, *D*. Because of the vanishing boundary condition on 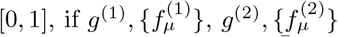 are all in ℋ_0_, so are *g*^*′*^, 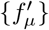, and thus 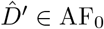. Therefore, AF_0_ is a vector subspace of 𝒞(ℝ ^*D*^, ℝ).

#### Elementary functions

Following the line of [94], we can build approximating functions for the indicator of half-planes in ℝ ^*D*^, i.e. 𝟙{***κ*** · ***d*** > *b*} for arbitrary ***d*** ∈ ℝ ^*D*^, *b* ∈ ℝ.

Let ℋ_0_ ∋ *T* : *z* 1 ↦ 2(1 − |2*z* − 1|) be the normalised triangle function over [0, 1]. Given ***d*** ∈ ℝ ^*D*^ and *b* ∈ ℝ, for *A*> 0, let ***f*** (*z*) = *AT* (*z*)***d***, *f*_0_(*z*) = −*AT* (*z*)*b* and *g*(*z*) = *T* (*z*) in Eq.(E.2). Then for *A* → ∞,

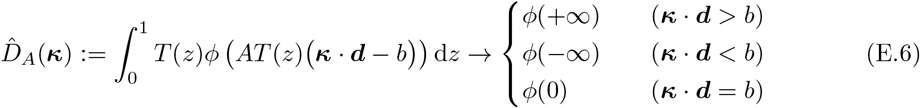

By assuming that *ϕ* is sigmoidal (i.e. *ϕ*(+ ∞) = 1, *ϕ*(− ∞) = 0), we see that 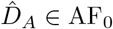 approximates the indicator of the half-plane in direction ***d***, given *A* large enough. The constant function 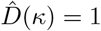 can also be trivially constructed.

Since linear combinations of indicators of half-planes are dense in *C*(ℝ ^*D*^, ℝ) with compact support [94], arbitrary functions *D*_*α*_ can be approximated by functions of the form of Eq.(E.2).

## Footnotes

1 so that the field of instantaneous firing rates, given by *ϕ*(*v*_*t*_(*z*)), is continuous over the similarity space.

